# A general role of zinc binding domain revealed by structures of σ^28^-dependent transcribing complexes

**DOI:** 10.1101/2020.01.02.893370

**Authors:** Wei Shi, Wei Zhou, Baoyue Zhang, Shaojia Huang, Yanan Jiang, Abigail Schammel, Yangbo Hu, Bin Liu

## Abstract

In bacteria, σ^28^ is the flagella-specific sigma factor that controls the expression of flagella-related genes involving bacterial motility and chemotaxis. However, its transcriptional mechanism remains largely unclear. Here we report cryo-EM structures of σ^28^-dependent transcribing complexes on a complete flagella-specific DNA promoter. The structures reveal how σ^28^-RNA polymerase (RNAP) recognizes promoter DNA through strong interaction with −10 element but weak contact with −35 element to initiate transcription. In addition, we observed a distinct architecture in which the β′ zinc binding domain (ZBD) of RNAP stretches out from its canonical position to interact with the upstream non-template strand. Further *in vitro* and *in vivo* assays demonstrate that this interaction facilitates the isomerization of RNAP-promoter closed to open complex due to compensating the weak interaction between σ4/−35 element, and suggest that ZBD-relocation is a general mechanism employed by the σ^70^-family factors to enhance transcription from promoters with weak σ4/−35 element interactions.

## Introduction

Multi-subunit DNA-dependent RNA polymerase (RNAP) is the core enzyme responsible for transcription, the first step of gene expression in cells. In bacteria, the RNAP core enzyme comprises four types of evolutionarily conserved subunits (α_2_ββ′ω) (Borukhov & Nudler, 2008; Murakami & Darst, 2003), which is unable to recognize specific promoter sequences alone. Transcription initiation is tightly regulated by sigma factors, in which the core enzyme first binds a specific sigma factor to form a holoenzyme and then recognizes specific promoters. The subsequent step is isomerizing from closed RNAP-promoter complex (RPc) to open RNAP-promoter complex (RPo) with melted base pairs, preparing to initiate transcription (Browning & Busby, 2016; Feklistov & Darst, 2011; Ruff et al, 2015; Saecker et al, 2011).

On the basis of the structural and functional differences, bacterial sigma factors could be classified into two main families: the primary σ^70^ factor family and the σ^54^ factor family for nitrogen regulation and some stress responses (Feklistov et al, 2014). The σ^70^ family factors are then subdivided into four major groups according to the different compositions of the conserved domains - σ1.1 (σR1.1 region), σ2 (σR1.2, σNCR, and σR2.1-2.4 regions), σ3 (σR3.0 and σR3.1 regions) and σ4 (σR4.1 and σR4.2 regions) (Feklistov et al, 2014; Paget, 2015). Sigma factors in group 1 (σ^70^ in *E. coli*) contain all the conserved domains, while group 2 sigma factors (σ^S^ or σ^38^ in *E. coli*) lack σNCR region, group 3 sigma factors (σ^28^ in *E. coli,* also known as RpoF or FliA) are short of σR1.1, σNCR and σR1.2 regions, and group 4 (extra cytoplasmic function, ECF) sigma factors are the most stripped-down version, possessing only two essential domains σ2 and σ4 (Osterberg et al, 2011; Paget, 2015).

As a representative group 3 sigma factor, σ^28^ controls flagellum biosynthesis in all motile Gram-negative and Gram-positive bacteria (Paget, 2015), and is indispensable for motile bacteria to compete with other microorganisms and survive in the adverse conditions like poor nutrition (Zhao et al, 2007). In the cytoplasm, σ^28^ factor is sequestered by an anti-sigma factor FlgM to form an inactive σ^28^/FlgM complex. Under inducible conditions, the cytoplasmic FlgM is transported out of the cell and the freed σ^28^ factor can bind to the core RNAP and activate the transcription of the flagellar genes (Gillen & Hughes, 1991; Hughes et al, 1993). The X-ray crystal structure of σ^28^/FlgM complex has revealed the inhibitory mechanism of FlgM in which FlgM wraps around outside and packs three conserved domains of σ^28^ together into a compact conformation, suggesting that σ^28^ must also undergo a large structural rearrangement to bind RNAP and form holoenzyme (Sorenson et al, 2004).

The well-studied transcription initiation complex (TIC) structures of group 1 sigma factors (Murakami et al, 2002b; Zhang et al, 2012; Zuo & Steitz, 2015) and the recent TIC structures of group 2 sigma factor (Liu et al, 2016) and group 4 sigma factors (Li et al, 2019; Lin et al, 2019) have greatly facilitated our understanding of how these sigma factors recognize respective promoter elements and initiate transcription. As the second richest sigma factor in *E. coli*, the level of σ^28^ is maintained at 50% of the level of σ^70^ factor (Jishage et al, 1996), and the ratio of σ^28^-RNAP holoenzyme to total holoenzymes is around 14%, only second to σ^70^-RNAP holoenzyme (Maeda et al, 2000). However, as yet, there is no structure of RNAP holoenzyme or transcription complex for σ^28^ to illustrate the mechanism of transcription initiation. In this study, we assembled the intact functional *E. coli* transcribing complex with the flagella-specific sigma factor - σ^28^, and determined the first TIC structures of group 3 factors at around 3.9 Å resolution. The structures show that σ^28^-RNAP has strong interaction with promoter −10 element but weak contact with −35 element. Intriguingly, the β′ zinc binding domain (ZBD), also previously known as zinc ribbon region (ZNR) (Lane & Darst, 2010), of RNAP stretches out from its canonical position to interact with non-template strand (NT-strand) in a distinct architecture, which consequently stabilizes promoter binding to advance transcription initiation by compensating the weak interaction between σ4/−35 element. Further *in vitro* and *in vivo* tests reveal that ZBD-relocation facilitates the isomerization from RNAP-promoter closed complex to open complex, and also suggest that the relocation of β′ ZBD is a general mechanism employed by sigma factors with weak σ4/−35 element interactions in transcription initiation. These observations advance our understanding of transcription initiation by RNAP from weakly-interacted or non-conserved promoters, the majority in bacteria (Ettwiller et al, 2016).

## Results

### Overall structure of the *E. coli* σ^28^-dependent transcribing complex

To obtain the cryo-EM structure of the *E. coli* σ^28^-dependent transcribing complex, we assembled the complex with RNAP core enzyme, σ^28^ factor, the synthetic DNA scaffold (from −39 to +15, positions relative to the transcription start site +1) (Fig 1A). To test RNA synthesis ability on the promoter by the σ^28^-RNAP holoenzyme, we also incubated the complex with nucleotides (ATP and CTP) and Mg^2+^, and isolated the target transcribing complexes (here we called TICs) from the reaction mixture (Fig EV 1A and B).

**Figure 1.**
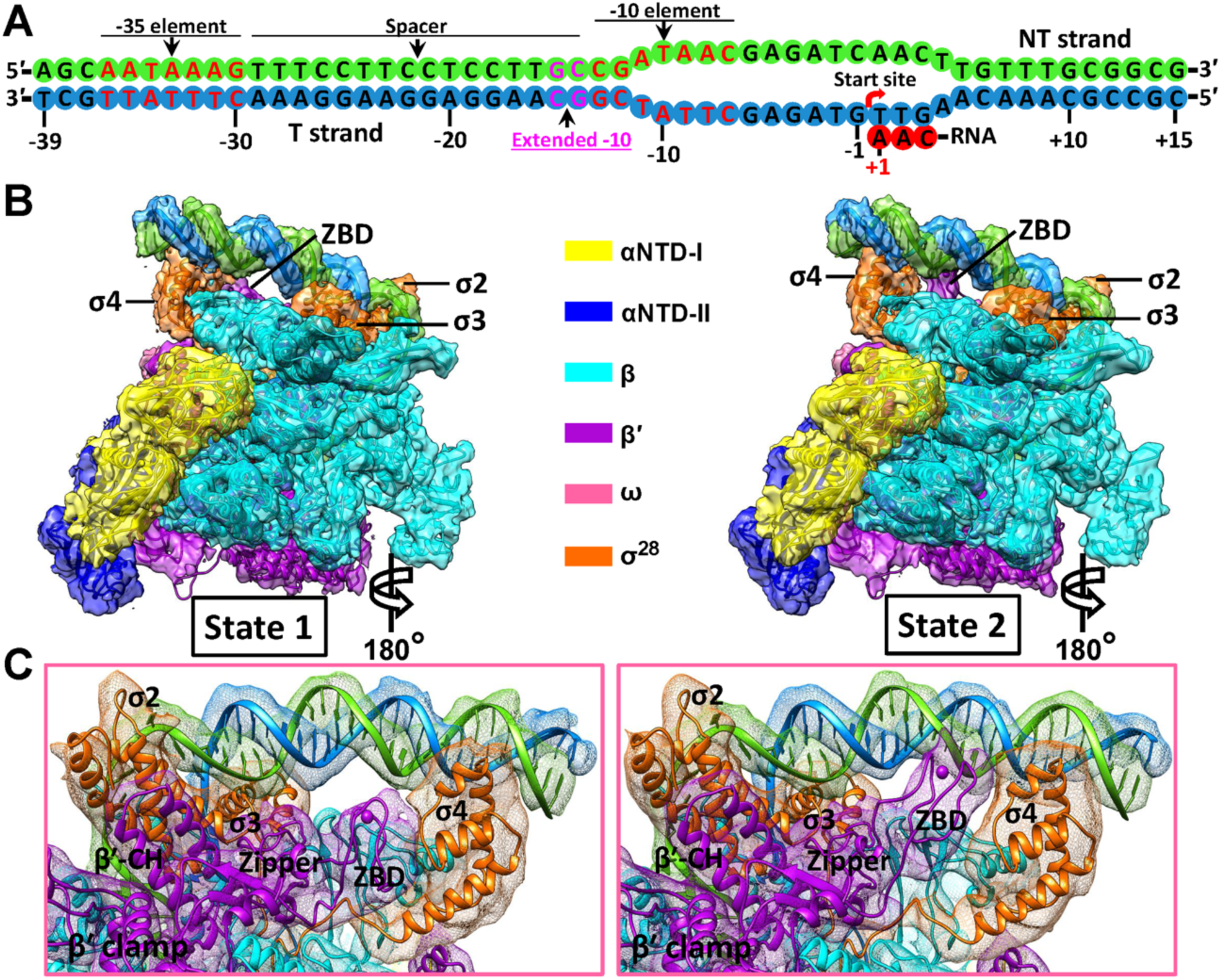
Cryo-EM reconstructions of the σ^28^-TICs at the two states. A. Schematic representation of the synthetic promoter DNA scaffold (54-bp) in the σ^28^-TICs. B. Overviews of the cryo-EM reconstruction maps of the *E. coli* σ^28^-TICs at the state 1 (left, 3.86 Å resolution) and state 2 (right, 3.91 Å resolution), respectively. The individually colored density maps, created by color zone and split in Chimera and shown in a contour of 8 root-mean-square (RMS), are displayed in transparent surface representation to allow visualization of all the components of the complex. C. Zoom-in views of β′ ZBDs in the σ^28^-TICs. The split density maps on the β′ subunits are displayed at 8 RMS (left) and 6 RMS (right), respectively. Others are same as in panel B.

The cryo-EM single-particle reconstruction of the complex showed two different architectures corresponding to two states (Fig 1B and C). Both state 1 and state 2 structures were reconstructed at the overall resolution of 3.86 Å and 3.91 Å respectively (Fig EV 1C-G, and Table EV1). The densities in the cryo-EM maps were well fit by the core RNAP, which shows a similar overall architecture to that observed in other TICs (Liu et al, 2017; Liu et al, 2016; Zuo & Steitz, 2015). The σ^28^ factor has relatively lower sequence identity (22.7%) and similarity (38.9%) to σ^70^ factor (Fig EV 2A). However, the folding pattern of σ^28^ factor in TICs shows high similarity with that of σ^38^ or σ^70^ throughout the conserved regions, from σR2.1 to σR4.2 regions (Fig EV 2B), indicating the conserved structures of the σ^70^ family in TICs. The structures of the two states also display a well-ordered density of nucleic acids including the bubble region and the newly-synthesized RNA (Fig EV 2C and D).

The two structures of σ^28^-TIC complexes show high similarity, with a root-mean-square deviation (RMSD) of 0.255 Å (Cα aligned). The major difference in two structures lies in the location of β′ ZBD of RNAP. In state 1 structure, ZBD is adjacent to the σ4 domain and does not interact with the upstream DNA; while in state 2 structure, ZBD gets close to the upstream DNA and makes contact with the NT-strand DNA (Fig 1B and C).

### Stable promoter recognition on the −10 element but weak on the −35 element

The σ^28^ factor recognizes the −10 and −35 elements via its σ2 and σ4 domains, respectively, which is similar to other σ^70^-family factors in TIC structures (Bae et al, 2015; Zuo & Steitz, 2015). The strong density on the −10 element suggests a stable recognition here. Five positively-charged arginine residues of σ^28^ have pairs of polar interactions with nucleotides of the template −10 element: R95/−13G, R95/−12C, R98/−12C, R98/−11T, R84/−12C, R34/−11T and R94/−11T. In addition, several residues in the β and β′ subunits (βN494, βK496, βR470, β′S319, and β′R259) also make contacts with the template strand (T-strand) −10 element via hydrogen bond or polar interactions (Fig 2A and B), which are similar to those observed in the σ^38^-TIC (Liu et al, 2016). The NT-strand −10 element is contacted by the region of σR2.3, and the path down to the main cleft is the same as that in previously reported TICs (Feklistov & Darst, 2011; Zhang et al, 2012). Multiple residues of the σR2 region contribute to the recognition of the sequence: R58/−13C, R74/−13C, Q73/−12G, Q63/−11A, T69/−9A and H26/−7C (Fig 2A and B). However, residues involved in recognizing the −10 element in σ^28^ are not conserved compared to those in σ^70^ and σ^38^ (Fig EV 2A), suggesting high specificity of σ^28^ in −10 recognition. The C_−13_G_−12_ATAAN_−7_ is conserved in promoters of σ^28^-dependent flagellar genes (Fig 2C and Table EV2) (Fitzgerald, 2014; Zhao et al, 2007), and the major difference with that of σ^70^ is the −13C and −12G. Mutation of either of the two nucleotides in a σ^28^-dependent promoter *tar*p (Koo et al, 2009) greatly decreased the promoter activity (Fig 2D). These *in vitro* data therefore show that σ^28^ factor specifically recognizes a subset of promoters with conserved −13C and −12G.

**Figure 2.**
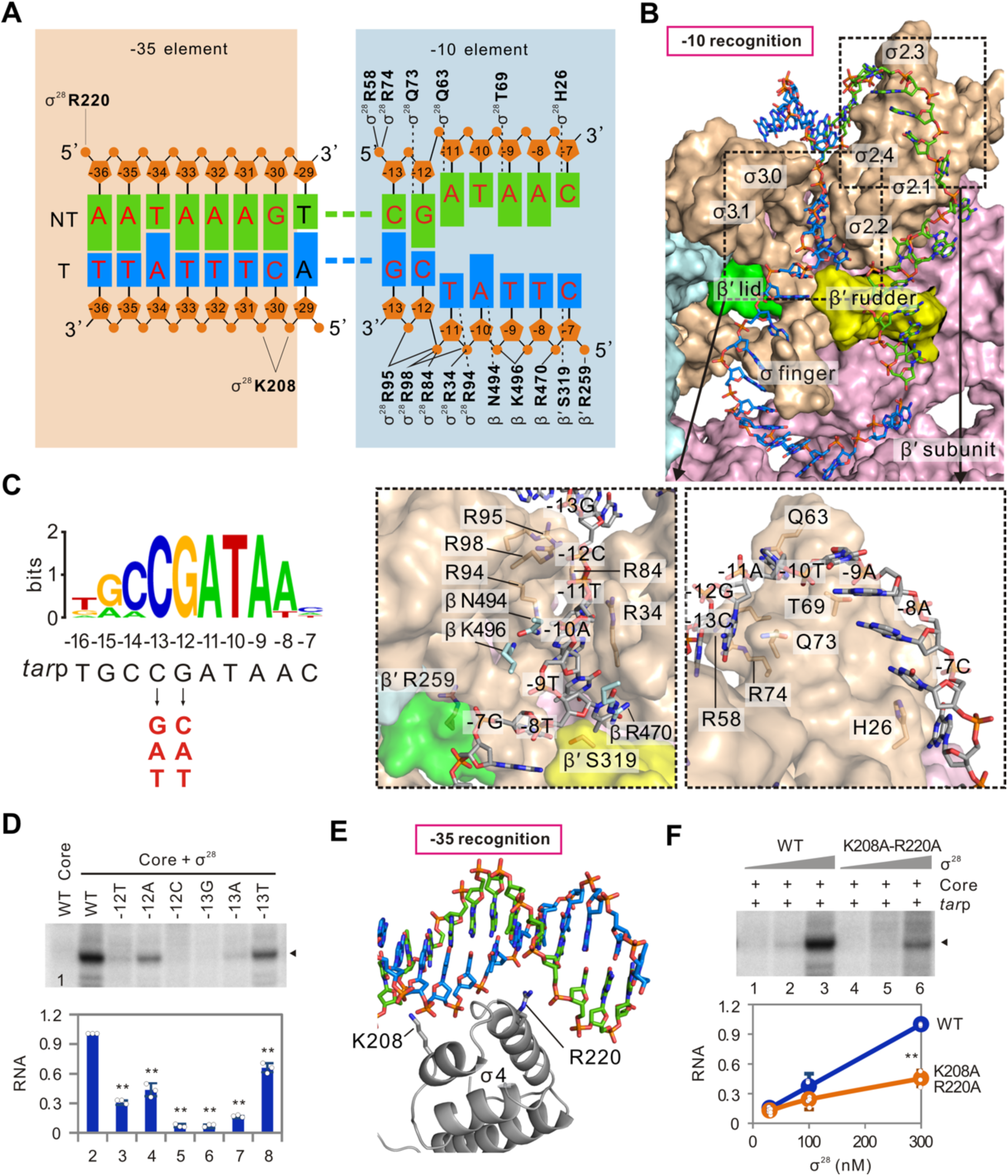
Promoter recognition by the σ^28^ factor. A. Summary of protein-nucleic acid interactions. Solid and dashed lines represent polar interactions and hydrogen bonds, respectively. The conserved −35 and −10 elements are colored in red letters. B. Recognition of the promoter −10 element. The RNAP is shown in surface representation: β subunit, cyan; β′ subunit, pink; β′ lid, green; β′ rubber, yellow; and σ^28^, wheat. The right and left side insets at the bottom part are the zoom-in views of the interactions on the NT-strand and T-strand, respectively. C. Conserved sequences around the −10 element of flagellar promoters (from −16 to −7 positions) generated by WebLogo (Crooks et al, 2004). Promoter sequences used in this analysis are shown in Table EV2. Corresponding sequence of *tar*p is shown below the logos. The mutated nucleotides in *tar*p are shown in red. D. Transcription of σ^28^-RNAP on wild-type or mutated *tar*p. Mutated nucleotides on non-template strand of *tar*p are shown. RNA products that are marked by a solid triangle were quantified and normalized to the signal obtained on the wild-type *tar*p (WT). Individual values of biological replicates (n=3) are shown as dots, and the means and standard deviations are displayed as error bars. ** p<0.01. E. Recognition of the promoter −35 element by the σ4 domain. F. Influence of mutating σ4 on promoter recognition. Quantifications are mean ± SD from three determinations. ** p<0.01.

Apparently, the interactions between σ^28^ and −35 element are not as extensive as the recognition on −10 element. Two positively charged residues recognize the phosphate atoms of the −35 element: K208/−30C (T-strand), K208/−31T (T-strand), and R220/−36A (NT-strand) (Fig 2A and E). Mutation of the two residues (K208A-R220A) significantly decreased the transcriptional activity of σ^28^-RNAP holoenzyme on *tar*p (Fig 2F). The number of residues recognizing −35 element in σ^28^ is obviously less than that in σ^70^ (Campbell et al, 2002), indicating the interaction might be less extensive than that for σ^70^. The first round of focused classification and refinement also showed that only 33.33% of the particles have clear densities on −35 element region (Fig EV 1D), suggesting a weak recognition. In addition, on the basis of the comparison with the σ^38^-TIC structure (PDB ID 5IPL) and σ^70^-TIC structure (PDB ID 4YLN), the position of σ4 helices in our structure are similar to those in the σ^38^-TIC, while they are around 2-3 Å farther away from the −35 element than the σ4 helices in σ^70^-TIC (Fig EV3), indicating that the recognition on the −35 element in the σ^28^-TIC is weaker than that in the σ^70^-TIC.

### Recognition of the upstream NT-strand DNA in the spacer by β′ ZBD

Remarkably, β′ ZBD of RNAP adopts an extended conformation in the state 2 structure compared with that in the state 1 structure (Fig 1C). The β′ ZBD locates at the N-terminal of β′ subunit, and four cysteines of ZBD are coordinated by one zinc ion (Fig 3A). The β′ ZBD in the state 1 architecture is around 12 Å away from the upstream DNA, and makes contact with the σ4 domain; while in the state 2 structure, ZBD is dissociated from the σ4 domain with a shift of ∼ 9 Å and reaches to the upstream NT-strand DNA (Figs 1C, 3A and EV 4A). In the σ^38^-TIC structure (PDB ID 5IPL) and σ^70^-TIC structure (PDB ID 4YLN), β′ ZBDs situate at similar positions as observed in the state 1 structure. Superimposition of them with our state 2 structure also shows ∼ 10 Å distance shift of Zn^2+^ (Fig EV 4B and C). The interface on β′ ZBD possesses a patch of positive charges (Fig EV 4D). Together with the highly positively-charged regions of the σ2, σ3 and σ4 domains (Fig EV 4D), we propose that interactions on β′ ZBD might contribute to a more stable promoter binding and more efficient transcription initiation.

**Figure 3.**
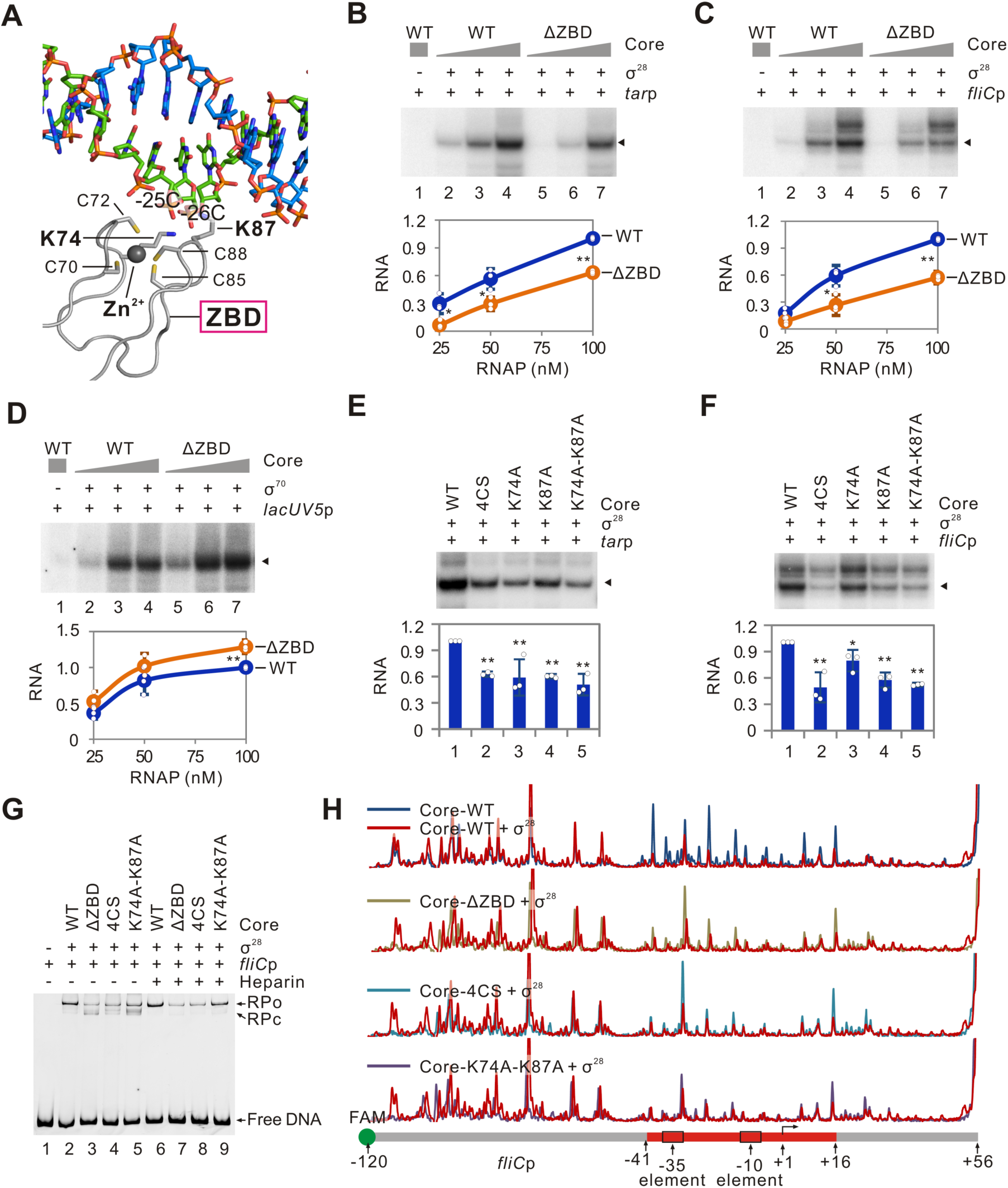
Role of β′ ZBD in activating σ^28^-dependent transcription. A. Specific interactions between the upstream DNA (−25 and −26 nucleotides) and β′ ZBD. B-D. Effects of deleting β′ ZBD on transcriptional activities of RNAP on different promoters. The *tar*p (**B**) and *fliC*p (**C**) were used as templates for σ^28^-dependent transcription, and the *lac*UV5p (**D**) was used in σ^70^-dependent transcription. RNA products indicated by solid triangles were quantified and the signal obtained from 100 nM wild-type RNAP was normalized to 1 in each test respectively. Data are mean ± SD from three determinations. Individual values of replicates are shown as dots. * p<0.05, ** p<0.01. E, F. Transcriptional activities of ZBD mutated σ^28^-RNAP on *tar*p (**E**) and *fliC*p (**F**). RNA products indicated by solid triangles were quantified from three determinations and are shown as mean ± SD. ** p<0.01. G. Bindings of σ^28^-RNAP with *fliC*p as detected by EMSA. The heparin-resistant RNAP-promoter open complex is indicated as “RPo”, and the heparin-sensitive RNAP-promoter closed complex is shown as “RPc”. H. DNase I footprinting of the *fliC*p complex with σ^28^-RNAP. Promoter DNA was labeled on the non-template strand. The promoter region protected by RNAP is indicated by a red box at the bottom panel.

Consistent with our hypothesis, comparing with wild-type RNAP, the ZBD-deleted RNAP (ΔZBD) showed reduced transcriptional activity when it was reconstituted with σ^28^ on *tar*p (Figs 3B and EV 5A) or *fliC*p promoter (Kundu et al, 1997) (Figs 3C and EV 5B), but did not decrease activity with the σ^70^ on the classical *lac*UV5p (Figs 3D and EV 5C), suggesting a specific role of β′ ZBD. Moreover, the ZBD interacts with the NT strand mainly through polar interactions: K74/−26C, K74/−25C and K87/−26C (Fig 3A). Single mutation of either K74 or K87 (named as K74A and K87A, respectively) or both residues (named as K74A-K87A), or mutation of four cysteines in β′ ZBD (named as 4CS) in the RNAP core enzyme all showed decreased transcriptional efficiency when reconstituted with σ^28^ (Figs 3E and F, and EV 5D and E), but not with σ^70^ on *lac*UV5p (Fig EV 5F). Additionally, the ZBD-mutated RNAPs showed decreased promoter binding (Fig 3G and H) and formed a higher ratio of unstable heparin-sensitive “closed complex” and a lower ratio of heparin-resistant “open complex” on *fliC*p when reconstituted with σ^28^ (Figs 3G and EV 5G), but did not significantly influence promoter binding with the σ^70^ on *lac*UV5p (Fig EV 5H and I). Together, these *in vitro* data reveal that β′ ZBD significantly contributes to promoter binding of σ^28^-RNAP by facilitating the isomerization of RNAP-promoter from “closed complex” to “open complex”.

### β′ ZBD contributes to expression of σ^28^-dependent genes and bacterial motility

To investigate if the roles of β′ ZBD revealed *in vitro* could be validated *in vivo*, we have tried to construct *E. coli* strains with mutations in the *rpoC* gene (encoding the β′ subunit) using a CRISPR/Cas9 system (Fig 4A) (Jiang et al, 2015). In spite of several attempts, we failed to obtain the strain with β′ ZBD deletion (see discussion). However, we have successfully obtained the mutants with single mutation at β′K74 (named as *Ec*-K74A) or β′K87 (named as *Ec*-K87A), and with double point mutations (named as *Ec*-K74A-K87A) (Fig 4A). Promoter activities of both *tar*p and *fliC*p, but not the native σ^70^-dependent *lac* promoter (*lac*p), were decreased in these mutants (Figs 4B and C, and EV 6A).

**Figure 4.**
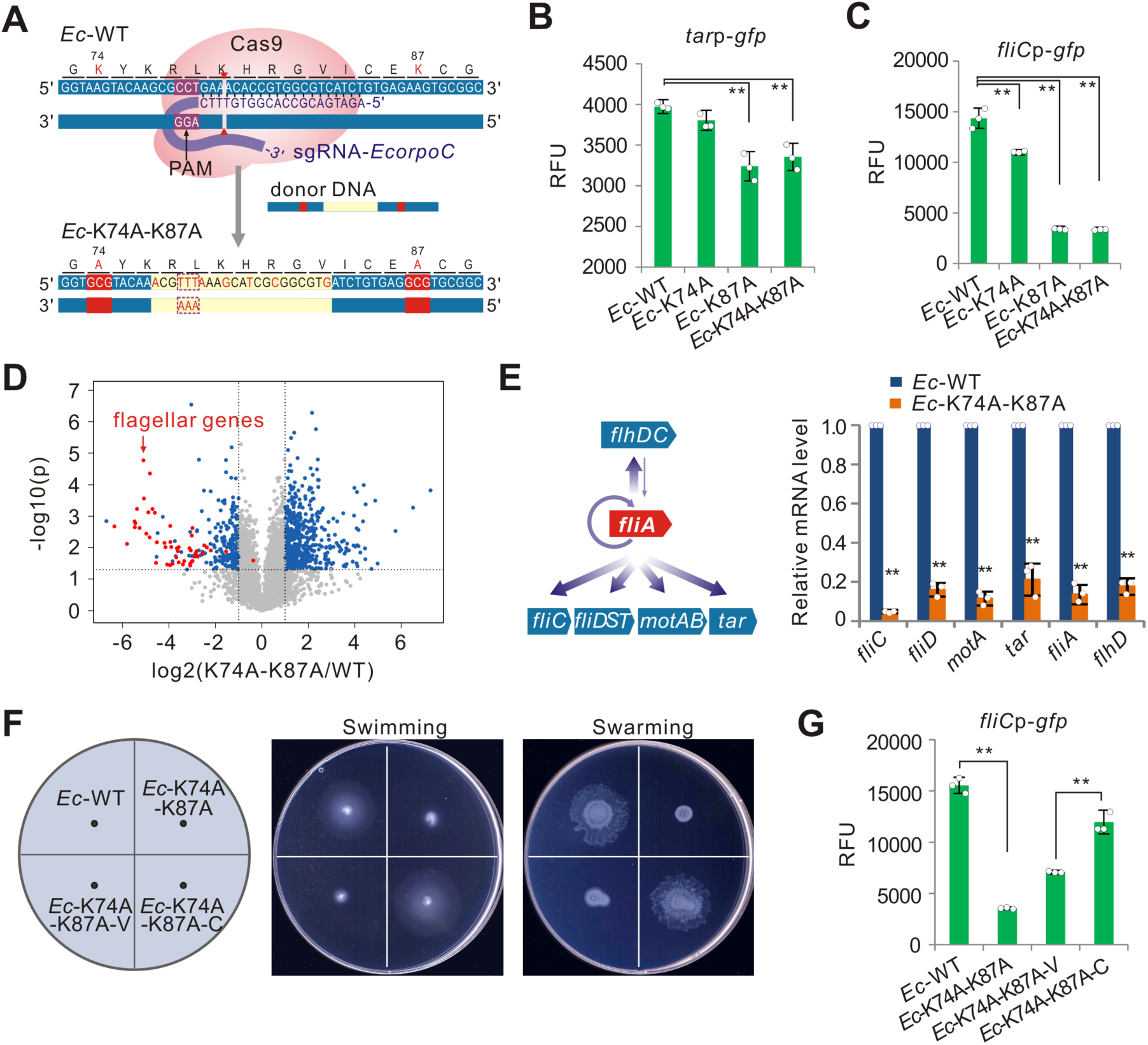
Importance of β′ ZBD in controlling transcription of σ^28^-dependent genes. A. Schematic diagram of constructing chromosomal mutations in *rpoC* gene in *E. coli* based on a CRIPSR/Cas9 system. A 20-nt guide RNA targeting a region between K74 and K87 on β′ subunit is shown in purple. Double-strand DNA fragments, which contains synonymous mutations (yellow background) in sgRNA targeting region to facilitate clone selection and aimed mutations at K74 or K87 or both residues (only example for constructing the *Ec*-K74A-K87A strain is shown), were used as donor templates. B, C. Promoter activities of *tar*p (**B**) and *fliC*p (**C**) in *E. coli* wild-type (*Ec*-WT) and its mutants expressing ZBD-mutated RNAP (*Ec*-K74A, *Ec*-K87A and *Ec*-K74A-K87A) shown as mean ± SD from three replicates. ** p<0.01. D. Transcriptional profiles of *Ec*-K74A-K87A and *Ec*-WT strains. Blue dots represent genes with significant difference between these two strains. Red dots represent flagellar genes. Data are shown as mean values of three independent colonies. E. Relative mRNA levels of flagellar genes in *Ec*-K74A-K87A strain comparing with that in *Ec*-WT (normalized to 1 for each gene). Diagram for the role of σ^28^ (*fliA* gene) in flagella regulon hierarchy (Clarke & Sperandio, 2005; Fitzgerald, 2014) is shown in the left panel. Data are mean ± SEM from three replicates. ** p<0.01. F. Motilities of *Ec*-K74A-K87A strains with over-expression of wild-type *rpoC* gene (*Ec*-K74A-K87A-C) or carrying the control plasmid (*Ec*-K74A-K87A-V). G. Activities of *fliC*p in *Ec*-K74A-K87A-C and *Ec*-K74A-K87A-V strains. L-arabinose was used at final concentration of 0.002%. Data are mean ± SD from three colonies. ** p<0.01.

Transcriptional profiling of *Ec*-WT and *Ec*-K74A-K87A strains showed that expression of all flagellar genes was decreased in the *Ec*-K74A-K87A strain (Fig 4D). Consistently, quantitate RT-PCR analysis displayed that transcription of all tested σ^28^-dependent flagellar genes (Fig 4E) presented decreased mRNA levels in the *Ec*-K74A-K87A strain. In addition, the *Ec*-K74A-K87A strain appeared almost non-motile in both swimming and swarming assays (Fig 4F), and mutation of either K87 or K74 also impaired bacterial motility (Fig EV 6B). Over-expression of σ^28^ (FliA) in the *Ec*-K74A-K87A strain could neither restore the *fliC*p activity nor the bacterial motility (Fig EV 6C and D), but both *fliC*p activity and bacterial motility can be partially restored by over-expression of a wild-type *rpoC* gene in the *Ec*-K74A-K87A strain (Fig 4F and G). These data suggest that the repressive effectiveness in the *Ec*-K74A-K87A strain is due to the mutation of ZBD but not repressed expression of σ^28^. Another constructed strain named *Ec*-CK, which carries all synonymous mutations in the *rpoC* gene of the *Ec*-K74A-K87A strain, behaved similarly as the *Ec*-WT strain (Fig EV 6E), further indicating that the loss of motility in the *Ec*-K74A-K87A strain is due to the mutations in K74 and K87 residues.

### β′ ZBD is generally important for transcription initiated by σ factors without strong σ4/−35 element interactions

Based on the fact that the interactions between σ2 and −10 element are strong but the ones between σ4 and −35 element are weak in σ^28^-TIC structures, and that β′ ZBD directly interacts with promoter region close to the −35 element to stabilize the RNAP-promoter complex, we propose that the contribution of β′ ZBD would compensate the weak interaction between σ4 with −35 element during transcription initiation. To test this hypothesis, we constructed a chimeric sigma factor, named as σ^28^-R4m, in which the σ4 region of σ^28^ was replaced by that of σ^70^ (Fig 5A). Correspondingly, the −35 element of σ^28^-dependent *tar*p promoter was also replaced by σ^70^-preferred “TTGACA” sequence to obtain a chimeric promoter *tar*p-35m (Fig 5A). The chimeric σ^28^-R4m efficiently recognized chimeric *tar*p-35m, and mutations in β′ ZBD did not significantly affect the transcriptional activity of σ^28^-R4m type RNAP on *tar*p-35m promoter (Fig 5B), supporting the hypothesis that the importance of β′ ZBD in transcription is associated with the strength of interaction between σ4 and −35 element.

**Figure 5.**
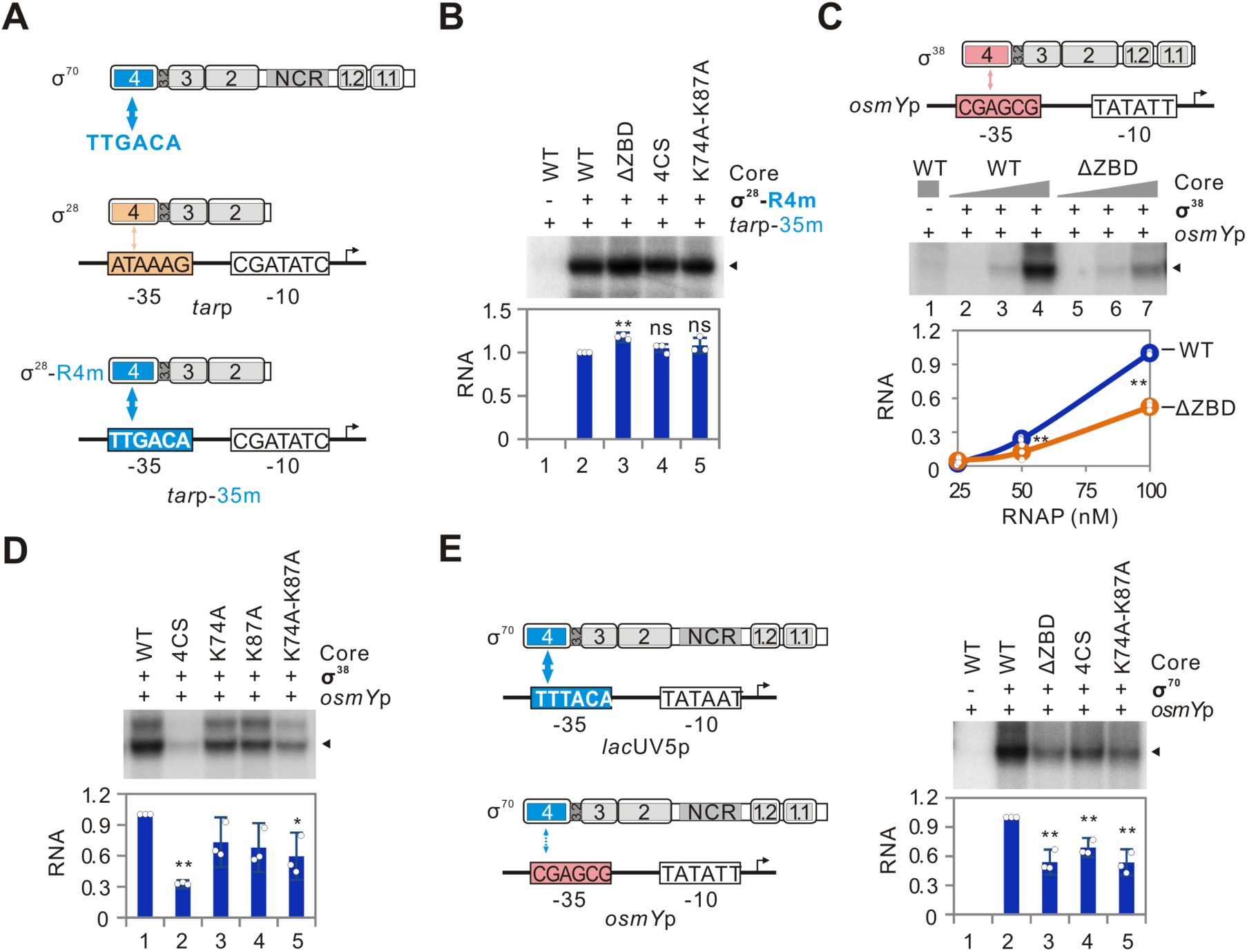
β′ ZBD contributes to transcription initiated by σ factors with weak σ4/−35 element interactions. A. Diagram for mutations in σ^28^ and *tar*p. Conserved domains in σ factor are shown as numbering. The region 4 of σ^28^ was replaced by that from σ^70^ to obtain the chimeric σ^28^-R4m, and the sequence of −35 element in *tar*p was replaced by “TTGACA” in the chimeric *tar*p-35m. The proposed strength for interaction between σ4/−35 in different σ/promoter pairs is indicated by the thickness of arrow. B. Activities of chimeric σ^28^-R4m in initiating transcription from *tar*p-35m, when reconstituted with wild-type (WT) or ZBD mutated RNAP core enzyme. Quantifications of RNA products indicated by solid triangles from three determinations are shown as mean ± SD at the bottom panel. ns, not significant; * p<0.05; ** p<0.01. C-E. Effects of mutating β′ ZBD on transcriptional activities of σ^38^-RNAP (**C**, **D**) or σ^70^-RNAP (**E**) on *osmY*p. Quantifications were performed from three determinations and are shown as mean ± SD. * p<0.05, ** p<0.01.

In addition to σ^28^, the σ^38^ is known to recognize promoters with high sequence variations at the −35 element (Maciag et al, 2011; Wise et al, 1996), suggesting a weak interaction between σ4 and −35 element. Intriguingly, mutations in ZBD also decreased the transcriptional efficiency of σ^38^-RNAP holoenzyme on a σ^38^-dependent *osmY*p promoter (Fig 5C and D), which contains a conserved −10 element and a non-conserved −35 element (Fig 5C) (Yim et al, 1994). Consistently, the activity of *osmY*p was decreased in *Ec-*K74A-K87A strain and partially restored in the *Ec*-K74A-K87A strain with over-expression of wild-type *rpoC* gene (Fig EV 6F).

The σ^70^ can also recognize promoters with non-conserved −35 element, such as *osmY*p (Colland et al, 2000). Comparing with the *lac*UV5p, the non-conserved −35 element in *osmY*p would decrease the strength of σ4/−35 element interaction for σ^70^-RNAP (Fig 5E). Mutations in β′ ZBD all decreased the activity of σ^70^-RNAP holoenzyme on *osmY*p (Fig 5E), which is different from the results on *lac*UV5p (Fig EV 5C and F). These data indicate that β′ ZBD is also important for σ^70^ to recognize promoters with non-conserved −35 element. Consistently, the *Ec*-K74A-K87A strain showed the delayed bacterial growth (Fig EV 6G), which could not be explained by functional deficiency of σ^28^ or σ^38^, since the strains with deletion of *fliA* gene (encoding the σ^28^) or *rpoS* gene (encoding the σ^38^) in *E. coli* did not affect bacterial growth (Schellhorn et al, 1998; Wood et al, 2006).

Altogether, both *in vitro* and *in vivo* data suggest that, independent of the types of sigma factors, strengthening the interaction between σ4 and −35 element would reduce the dependence of β′ ZBD domain in transcription, but transcription from promoters with weak interaction between σ4 and −35 element requires the contribution of β′ ZBD domain for maximal efficiency.

## Discussion

In bacterial RNA polymerase, the ZBD located at the N-terminal portion of RNAP β′ subunit. ZBD has been implicated in protein-DNA interactions at the downstream of catalytic center to stabilize the transcription elongation complex (Nudler et al, 1998), but it is reported later that ZBD interacts with product RNA located upstream of the catalytic center and the RNA-DNA hybrid in the elongation complex (King et al, 2004). It was also reported that ZBD participates in the salt-resistant interaction with double-stranded DNA and the Cys mutation renders the elongation complex extremely sensitive to salt and considerably less processive (Nudler et al, 1996). As for the transcription initiation, the mycobacterial TIC with RbpA (a transcription regulator in mycobacteria) shows that the core domain of RbpA makes extensive contacts with β′ ZBD domain(Hubin et al, 2017a), indicating a possible role of ZBD in transcription initiation.

Here, we provide first evidence for the role of β′ ZBD in transcription initiation. During the process of transcription initiation, β′ ZBD in core RNAP is first in a canonical position without extension or relocation. The unconstrained σ^28^ factor binds the core RNAP to form σ^28^-RNAP holoenzyme, which binds to the specific promoter to form an RPc. The unstable RPc can isomerize into a stable RPo through formation of several intermediate complexes (Boyaci et al, 2019; Saecker et al, 2011). The isomerization process requires stable RNAP-promoter interaction (Saecker et al, 2011), but the interaction between σ4 and −35 element is relatively weak for σ^28^. Therefore, we propose that the relocation of ZBD results in the direct interaction with the NT-strand of promoter spacer region and an increase in efficiency of the promoter isomerization (Fig 6), which is supported by our EMSA data that mutation of ZBD increased the ratio of RPc/RPo (Fig 3G). Since the RPc is unstable, the total amount of RNAP-promoter complexes was decreased for ZBD-mutated RNAPs as observed in DNase footprinting assays (Fig 3H). Moreover, previous studies have suggested that the intrinsic zinc ions in RNAP might play regulatory roles in substrate or template binding (Wu & Wu, 1987) and initiation of RNA synthesis (Scrutton et al, 1971). Additionally, Murakami *et al*. also proposed the possibility of the interaction of β′ ZBD with DNA backbone in the spacer between −22 on the T strand and −27 on the NT strand based on the structure of a RNAP holoenzyme-DNA complex (Murakami et al, 2002a), but they did not suggest any function of β′ ZBD. Our structures and biochemical assays demonstrated the positive role of ZBD in controlling transcription initiation and provided important and direct evidences to support these previous biochemical studies and proposals.

**Figure 6.**
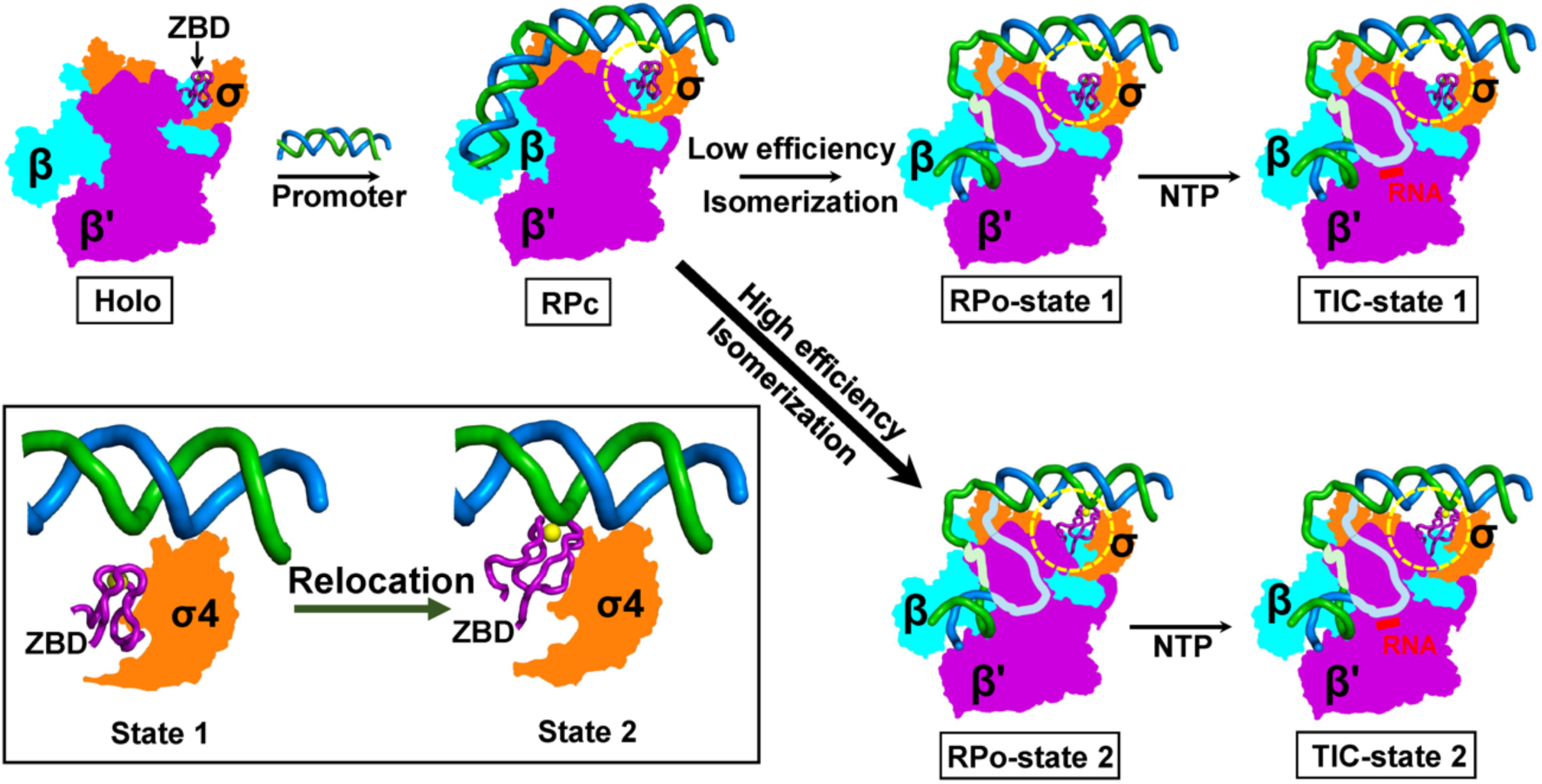
The ZBD-relocation mechanism in transcription initiation. A schematic cartoon model of ZBD-relocation mechanism during σ^28^-mediated transcription initiation is presented. The ZBD-relocation facilitates the isomerization of closed RNAP-promoter complex (RPc) to open RNAP-promoter complex (RPo) by contributing to RNAP-promoter interaction. The RPc with the relocated ZBD (state 2) has a higher efficiency in isomerizing to RPo than the one with the state 1 ZBD, consequently advancing transcription efficiency. The inset is the zoom-in view of ZBD relocation from state 1 to state 2.

The novel role of β′ ZBD is associated with the strength of interaction between σ4 and −35 element. The β′ ZBD has positive effects in σ^28^-, σ^38^- and even in some σ^70^-directed transcription, but not in σ^70^-directed transcription on strong promoter, which may be due to a strong σ4/−35 element recognition in the σ^70^-TIC is sufficient to stabilize the whole upstream DNA for isomerization and transcription initiation. Promoter sequence analysis based on identified transcriptional start sites in *E. coli* (as well as in other bacteria) showed that most of promoters contain conserved −10 element but not conserved −35 element (Ettwiller et al, 2016; Hubin et al, 2017b), which suggests that β′ ZBD may be important for transcription of those genes. In support of this hypothesis, the K74A-K87A mutation in ZBD altered expression of a quarter of genes in *E. coli* (Fig 4D). The analyses may explain why we could not obtain the *E. coli* mutant with deletion of β′ ZBD. Thus, on the basis of our results and the previous reports, we suggest that the ZBD relocation is a general mechanism, by which bacterial RNAPs could enhance transcription initiation from promoters with weak σ4/−35 interactions.

With the nucleic-acid-binding property, the positively-charged residues of β′ ZBD make contacts only with the phosphate backbone of the upstream NT-strand DNA, and therefore should have no specific sequence requirement, which is different from sequence specific interaction between the β′ zipper domain with promoter spacer DNA (Yuzenkova et al, 2011). After analyzing the *E. coli* σ^70^-TIC (PDB ID 4YLN) (Zuo & Steitz, 2015), σ^38^-TIC (PDB ID 5IPL) (Liu et al, 2016) and *Mycobacterium tuberculosis* σ^H^-TIC (PDB ID 5ZX2) (Li et al, 2019) structures, we observed that the β′ ZBD domains have similar polar interactions with the upstream NT-strand DNA: K74 in σ^70^-TIC, K87 in σ^38^-TIC, and R77 in σ^H^-TIC (corresponding to K87 in *E. coli*). All these suggest that the two residues (K74 and K87) observed in our σ^28^-TIC structures, which are involved in stabilizing the promoter binding, are the common residues among these structures, indicating a general mode of action for β′ ZBD in transcription.

Sigma factors are recruited to form RNAP holoenzyme to recognize specific promoters and regulate transcription initiation. Previous structural studies have mainly focused on molecular mechanisms for transcription from promoters with strong interaction to RNAP (Liu et al, 2017; Murakami et al, 2002a; Zhang et al, 2012; Zuo & Steitz, 2015). In this study, the flagella-specific σ^28^-TIC structures provide key insights into mechanisms of transcription initiation from promoters with weak interaction between σ4 and promoter −35 element, in which the β′ ZBD is relocated to contact with the upstream NT-strand DNA in the spacer region to compensate the weak σ4/−35 element interaction and promote the isomerization of RNAP-promoter closed to open complex. Importantly, with the structural, the *in vitro* and *in vivo* assays combined together, the results show that ZBD-relocation is a general mechanism employed by a broader set of sigma factors/promoters in transcription initiation.

## Materials and Methods

### Bacterial strains, plasmids and oligonucleotides

Bacterial strains, plasmids and oligonucleotides generated and used in this work are listed in Tables EV3 and EV4.

### Expression and purification of RNAP core enzyme

The *E. coli* RNAP core enzyme was prepared by co-expression of genes for RNAP β subunit, C-terminally His-tagged β′ subunit, α subunit and ω subunit using pVS10-RNAP in *E. coli* BL21(DE3) as described previously (Belogurov et al, 2007; Liu et al, 2017; Liu et al, 2016; Zhi et al, 2003). Mutations in *rpoC* gene in pVS10-RNAP plasmid were introduced by overlap PCR. Mutated fragments were digested by *Sbf* I and *Hind* III, which were then inserted into the same digested pVS10-RNAP plasmid.

### Expression and purification of σ factors

For construction of the plasmid pET21a-Ecoσ^28^, the *E. coli* σ^28^ gene was PCR-amplified from *E. coli* genomic DNA and ligated into pET21a vector which contains C-terminal His_6_-tag using *Nde* I and *Xho* I restriction sites. The amplified *E. coli* σ^70^ gene was inserted into pET28a vector between *Nhe* I and *Hind* III restriction sites to obtain pET28a-EcorpoD, which expresses His_6_-tag at the N-terminal of σ^70^. The *E. coli* σ^38^ gene was also cloned into pET28a using a ClonExpress II One Step Cloning Kit (Vazyme). A fragment expressing His_6_-tagged MBP and a TEV cleavage site was introduced in fusion at the N-terminal of σ^38^ gene. All mutations in plasmids were introduced by oligos and the PCR-amplified plasmid fragments were then cycled using the ClonExpress II One Step Cloning Kit.

All these constructs were transformed into *E. coli* BL21(DE3) competent cells. The cells were grown in LB medium with 100 μg/mL ampicillin (pET21a) or 50 μg/mL kanamycin (pET28a) at 37 °C to an OD_600_ of 0.6, and induced with 0.5 mM isopropyl-β-D-thiogalactopyranoside (IPTG) to express σ factors at 16 °C for 16-18 h. The bacteria were then harvested by centrifugation, resuspended in lysis buffer containing 50 mM Tris-HCl, pH 8.0, 300 mM NaCl, 10 mM imidazole, 5% (v/v) glycerol, lysed by ultrasonication and then centrifuged at 80,000 g for 1 h using a T29-8×50 fixed angle rotor (Sorvall LYNX 6000 superspeed centrifuge, Thermo Scientific). For σ^28^ purification, the recombinant protein was purified through 5 mL HisTrap HP column (GE Healthcare), and the eluted protein was further purified through 5 mL HiTrap Heparin HP column (GE Healthcare). Finally, the protein was applied to a gel filtration column, 120 mL HiLoad 16/600 Superdex 200 pg (GE Healthcare) in a buffer containing 20 mM Tris-HCl, pH 8.0, 150 mM NaCl. The peak fractions were pooled and concentrated to around 5 mg/mL. The final sample was aliquoted, flash-frozen in liquid nitrogen, and stored at −80 °C until use. The recombinant σ^70^ was purified in a similar way, but a Superdex 200 Increase 10/300 GL column (GE Healthcare) was used for gel filtration analysis. The recombinant σ^38^ was firstly passed through 5 mL HisTrap HP column (GE Healthcare), and the eluted protein was diluted and incubated with purified His_6_-TEV protein expressed by pRK793 plasmid (Kapust et al, 2001). The σ^38^ protein was then passed through the HisTrap HP column again to remove His_6_-MBP and His_6_-TEV enzyme. Fractions passed through the HisTrap HP column were collected for further purification with the Superdex 200 Increase 10/300 GL column (GE Healthcare).

### Assembly and purification of σ^28^-TIC

The synthetic DNA scaffold used in assembly is the σ^28^-specific *tar*p promoter with minor changes in the sequence to form the bubble, which corresponds to the promoter region between position −39 and +15 relative to the transcription start site (Fig 1A). The promoter DNA was prepared by annealing the NT-strand DNA to an equal molar amount of T-strand DNA. The σ^28^-TIC was assembled by directly incubating RNAP core enzyme with a threefold molar amount of purified σ^28^ protein and the preformed promoter DNA scaffold in buffer A containing 20 mM Tris-HCl, pH 7.5, 50 mM NaCl, 5 mM MgCl_2_ at 37 °C for 15 min in the presence of ATP and CTP (0.2 mM each). The reaction mixture was then used for purification through size-exclusion chromatography with Superdex 200 Increase 10/300 GL column (GE Healthcare) in buffer A to remove extra σ^28^ protein and nucleic acids.

### Cryo-EM grid preparation and data acquisition

A drop of 3.5 μl of the purified σ^28^-TIC at about 1 μM was applied on Quantifoil R2/2 200 mesh Cu grids (EM Sciences) glow-discharged at 15 mA for 60 sec. The grid was then blotted for 3 sec at 4 °C under the condition of 100% chamber humidity, and plunge-frozen in liquid ethane using a Vitrobot mark IV (FEI). The grids were imaged using a 300 keV Titan Krios microscope (FEI) equipped with a Falcon III direct election detector (FEI) at the Hormel Institute, University of Minnesota. Data were collected in counted mode with a pixel size of 0.9 Å and a defocus range from −1.6 to −2.6 μm using EPU (FEI). Each micrograph consists of 30 dose-framed fractions, and was recorded with a dose rate of 0.8 e^-^/pixel/sec (1 e^-^/Å^2^/sec). Each fraction was exposed for 1 sec, resulting in a total exposure time of 30 sec and the total dose of 30 e^-^/Å^2^.

### Image processing

Cryo-EM data were processed using cisTEM (Grant et al, 2018), and the procedure is outlined in Fig EV1. A total of 3,993 movies were collected. Beam-induced motion and physical drift were corrected followed by dose-weighting using the Unblur algorithm (Grant & Grigorieff, 2015). The contrast transfer functions (CTFs) of the summed micrographs were determined using CTFFIND4 (Rohou & Grigorieff, 2015). From the summed images, particles were then automatically picked based on a matched filter blob approach with the parameters: maximum particle radius (80 Å), characteristic particle radius (60 Å), threshold peak height (1.5, standard deviation above noise), 30 Å of highest resolution was used, and high variance area and areas with abnormal local mean were avoided (Sigworth, 2004). In total, 1,067,697 particles were picked and selected to construct a refinement package with 160 Å of estimated largest dimension/diameter and 324 pixels of box size for particles. 2D classifications (Scheres et al, 2005) were performed using 300-40/8 Å (start/finish, high-resolution limit) data and without inputting starting reference. In the first round of 2D classification, 126 of 200 classes were manually chosen by removing poorly populated classes containing unsuitable particles and obvious Einstein-from-noise to construct a new refinement package of 760,853 particles, and subjected to the second round of 2D classification. Then, 46 of 200 classes were selected to construct a new refinement package with 435,521 particles, and subjected to Ab-Initio 3D reconstruction (Grigorieff, 2016) to generate an initial 3D model using 20-8 Å resolution data. The initial 3D model was set as the starting reference for further 3D Auto-refinement. Fourier Shell Correlation (FSC) at the criteria of 0.143 resulted in a 3.53 Å resolution for the map outputted from Auto-refinement (Chen et al, 2013). Due to the flexibility of upstream DNA, we performed a focused classification (4 classes) and refinement in 3D manual and local refinement using 300-8 Å resolution data and a sphere of 40 Å radius centered at the region of σ4. Good classes were obtained after 30 cycles of refinement. The third class (33.33%) was reconstructed with 145,159 particles to generate the architecture at 3.81 Å resolution, with a better density of upstream DNA with clear density on the −35 element. We further performed a second round of focused classification (4 classes) and refinement in 3D manual and local refinement with a sphere of 25 Å radius centered at β′ ZBD. Good classes were obtained after 30 cycles of refinement. The first class (30.20%) and the third class (21.60%) were reconstructed with 43,838 and 31,354 particles to generate two structures both at 3.91 Å resolutions: structure 1 (state 1) and structure 2 (state 2), which then were further refined to 3.86 Å and 3.91 Å resolution respectively. The other two classes (class 2 and class 4) have poor density on either the σ4 domain or β′ ZBD.

### Model building and refinement

The initial models were generated by docking the previous structures of the components in the RNAP core (PDB ID 6B6H) into the individual cryo-EM density maps outputted from the focused classification and refinement using Chimera (Pettersen et al, 2004) and COOT (Emsley & Cowtan, 2004). The 3.86 Å or 3.91 Å cryo-EM density maps allowed us to build the σ^28^ factor by first manually docking and then mutating the structure of σ^38^ (PDB ID 5IPL) according to the sequence alignment result and the density map, and to dock or build β′ ZBD and the RNA transcript (AAC) at the active site starting from +1 position in COOT. The clear corresponding density allowed us to manually build the promoter DNA scaffold in COOT as well. The ω subunits and β′ rim helices in two structures were also fitted in maps although poor densities suggest the flexibility or low occupancy, while no density allowed us to fit the CTD of the α subunit of RNAP into the map. Although the low occupancy of the ω subunit was observed in the structures, it should have no influence on the normal architecture of RNAP and β′ ZBD relocation since the main part of RNAP architecture is same as previously and the ω subunit is at least 50 Å apart from β′ ZBD. The intact models were then real-space refined using Phenix. The final maps were put into a P1 unit cell (a = b = c = 291.6 Å; α = β = γ = 90°) and the structural factors were calculated in Phenix (Adams et al, 2010). In the real-space refinement, minimization global and local grid search were performed with the secondary structure, rotamer, and Ramachandran restrains applied throughout the entire refinement. The final models have good stereochemistry by evaluation in MolProbity (Chen et al, 2010). The local resolution maps were estimated and generated by ResMap (Kucukelbir et al, 2014). The statistics of cryo-EM data collection, 3D reconstruction and model refinement were shown in Table EV1. All figures were created using Chimera (Pettersen et al, 2004) or PyMOL (Schrödinger LLC, https://pymol.org/2/).

### *In vitro* transcription assay

*In vitro* transcription assays were performed as previously described (Hu et al, 2012) with minor modifications. Briefly, promoter fragments (∼180 bp in length, shown in Table EV2) were amplified from *E. coli* genomic DNA and purified by a Gel Extraction Kit (Omega). RNAP holoenzyme was assembled by mixing 100 nM RNAP core and 300 nM σ factor (except those indicated in figures) in transcription buffer (20 mM Tris-HCl, pH 7.9, 50 mM NaCl, 5 mM MgSO_4_, 1 mM DTT, 0.1 mM EDTA, 5% glycerol). A promoter fragment (20 nM) was incubated with RNAP holoenzyme at 37 °C for 10 min. Transcription was initiated by the addition of 50 μM CTP, UTP and ATP, 5 μM GTP, and 1 μCi of [α-^32^P]GTP. The reactions were carried out at 37 °C for 10 min, and then stopped by 1 volume of 95% formamide solution. RNA products were heated at 70 °C for 5 min and then analyzed on denaturing (7 M urea) 16% polyacrylamide gel electrophoresis (PAGE).

### DNA binding analysis

For the electrophoretic mobility shift assays (EMSA), the fluorescein-labeled promoter fragments were mixed with RNAP holoenzyme as described in the *in vitro* transcription assay, and reactions were incubated for 10 min at 37 °C. Then, 10 μg/mL of poly(dA-dT) was added and incubated for 5 min at 37 °C. Afterwards, samples were loaded on 6% native 0.5×TBE-PAGE. Gels were scanned by Amersham Typhoon scanner (GE Healthcare). Where indicated, 10 μg/mL of heparin was added into samples before being loaded to the PAGE.

For DNase I footprinting analyses, the fluorescein-labeled promoter fragments were incubated with RNAP in a similar way as described in EMSA. After incubation of RNAP holoenzyme with promoter fragment for 15 min at 37 °C, samples were treated with 2 U/mL DNase I (Promega) for 1 min at 37 °C. Reactions were stopped by the addition of 10 mM EDTA (pH 8.0) and heated at 70 °C for 5 min. Samples were then loaded into an Applied Biosystems 3730xl DNA analyzer (Hu et al, 2012; Zhu et al, 2017).

### *E. coli* mutant construction

A CRISPR-Cas9 system (Jiang et al, 2015) was applied to construct the mutants based on *E. coli* MG1655 strain. Briefly, a small guide RNA (sgRNA) targeting *rpoC* gene (shown in Fig 4A) was introduced in pTargetF plasmid using the ClonExpress II One Step Cloning Kit to obtain pTargetF-EcorpoC. The MG1655 strain was first transformed with a Cas9 expressing plasmid pCas. Afterward, the pCas containing MG1655 cells were co-transformed with the pTargetF-EcorpoC plasmid and an 1168 bp donor DNA (containing synonymous mutations in sgRNA targeting region and mutations at K74A, or K87A, or both where needed). Mutants were confirmed by DNA sequencing. The plasmids in these strains were cured as described previously (Jiang et al, 2015). The final mutants were named as *Ec*-K74A, *Ec*-K87A and *Ec*-K74A-K87A, respectively. The strain contains only synonymous mutations was named as *Ec*-CK, and the parent strain was renamed as *Ec*-WT.

For constructing the complementary strains, wild-type *rpoC* gene was amplified from *E. coli* genomic DNA and cloned into pBAD22 plasmid (Guzman et al, 1995) using the ClonExpress II One Step Cloning Kit. The obtained plasmid was named as pBAD-EcorpoC, and was transformed into *Ec*-K74A-K87A strain to get strains named as *Ec*-K74A-K87A-C. The *Ec*-K74A-K87A strain transformed with pBAD22 plasmid was named as *Ec*-K74A-K87A-V. Similar procedure was applied for constructing pBAD-FliA plasmid.

### RNA extraction, RNA-seq and qRT-PCR analyses

For analyzing expression of flagellar genes, *E. coli* MG1655 and its mutants were grown in LB medium at 30 °C to late logarithmic growth phase (OD_600_ 0.9-1.0). RNA was extracted using TRIzol reagent (Invitrogen) as described by manufacturer’s protocol. For RNA-seq analysis, the rRNA in extracted RNA was removed by Ribo-off rRNA Depletion Kit (Vazyme), RNA library was constructed using a NEBNext^®^ Ultra™ Directional RNA Library Prep Kit for Illumina (NEB) and sequenced by Illumina HiSeq X Ten platform.

For quantitative RT-PCR (qRT-PCR) analysis, RNA was first reverse transcribed to cDNA using M-MLV (Promega). The relative amount of target mRNA was analyzed by quantitative RT-PCR (qRT-PCR) using an iTaq Universal SYBR Green Supermix (Bio-Rad) following the manufacturer’s instructions. Relative transcriptional levels of tested genes were normalized to the 16S rRNA levels.

### *In vivo* promoter activity test and motility assay

*In vivo* promoter activity was tested using a *gfp* reporter fusion plasmid named as pGT, which was constructed by integration of three fragments: p15A ori, kanamycin resistance gene, and a promoter-less *gfp* fragment. Promoter fragments were cloned into pGT using the ClonExpress II One Step Cloning Kit. The pGT constructs were transformed into *E. coli* MG1655 and its derivative strains, which were grown at 30 °C to late logarithmic growth phase. The fluorescence intensity was quantified by Synergy HT plate reader (Bio-Tek) in 96-well black plate. The promoter activities are shown as relative fluorescence units (RFU), determined as the fluorescence intensities per OD_600_. The native activity of *lac* promoter in each strain (cells were grown at 30 °C in the presence of 0.2 mM IPTG) was tested by measuring the β-galactosidase expression level (Hu et al, 2009).

For swimming motility assay, 2 μl of each strain (30 °C, late logarithmic growth phase) was stabbed into semi-solid agar medium (1% tryptone, 0.5% NaCl, 0.3% Difco agar) (Burkart et al, 1998). Plates were incubated at 24 °C for 14 h for imaging. For swarming test, 2 μl of each strain was spotted on LB plate containing 0.6% Difco agar and 0.5% glucose (Burkart et al, 1998), which was then incubated at 30 °C for 15 h before imaging. L-arabinose was added at 0.2% into plates for strains carrying pBAD22, pBAD-EcorpoC or pBAD-FliA plasmid.

### Quantification and statistical analysis

Quantification, statistical analysis, and validation are implemented in the software packages used for 3D reconstruction and model refinement. RNAs from three replicates of *in vitro* transcription assays and shifted DNAs in EMSA assays were quantified by ImageJ software. Statistical analyses were performed using the unpaired Student’s *t*-test (two-tailed) between each of two groups. Significant of difference in RNA-seq analysis was computed using R (version 3.2.2).

## Data and Software Availability

The accession numbers of the 3.86 Å resolution EM map of σ^28^-TIC at the state 1, and the 3.91 Å resolution EM map of σ^28^-TIC at the state 2 and their corresponding coordinates reported in this paper are EMDB: 20394 and 20395, and PDB: 6PMI and 6PMJ, respectively. The accession number for the sequencing data reported in this study is SRA: SRP239152.

## Acknowledgments

The cryo-EM data were collected at the cryo-electron microscopy facility in the Hormel Institute, University of Minnesota, which is funded by the Hormel Foundation. This work was supported by the starting-up funding granted to B.L. from the Hormel Institute, University of Minnesota and the National Natural Science Foundation of China to Y.H. (#31670134). Support from the Youth Innovation Promotion Association CAS to Y.H. and help from the Core Facility and Technical Support of Wuhan Institute of Virology in radioactive and fluorescent tests are also acknowledged.

## Author contributions

W.S., W.Z., Y.J. and A.S. performed molecular cloning and protein sample preparations. W.S. assembled complexes for structure determination. W.S. and B.L. performed cryo-EM grid preparation, screening, and optimization. B.L. conducted high throughput data collection on Titan Krios. W.S. and B.L. performed image processing, map generation, model building and refinement, and structural analysis. W.Z. and Y.H. performed *in vitro* transcription and DNA binding analyses. B.Z. and Y.H. constructed *E. coli* mutants and performed most of *in vivo* tests. S.H., W.Z. and Y.H. extracted RNA and performed RNA-seq and qRT-PCR analyses. W.S., Y.H. and B.L. wrote the manuscript with contributions from all authors.

