## Supplementary Figures and Tables for "A general role of zinc binding domain revealed by structures of σ^28^-dependent transcribing complexes"

### Expanded View Figures and Tables

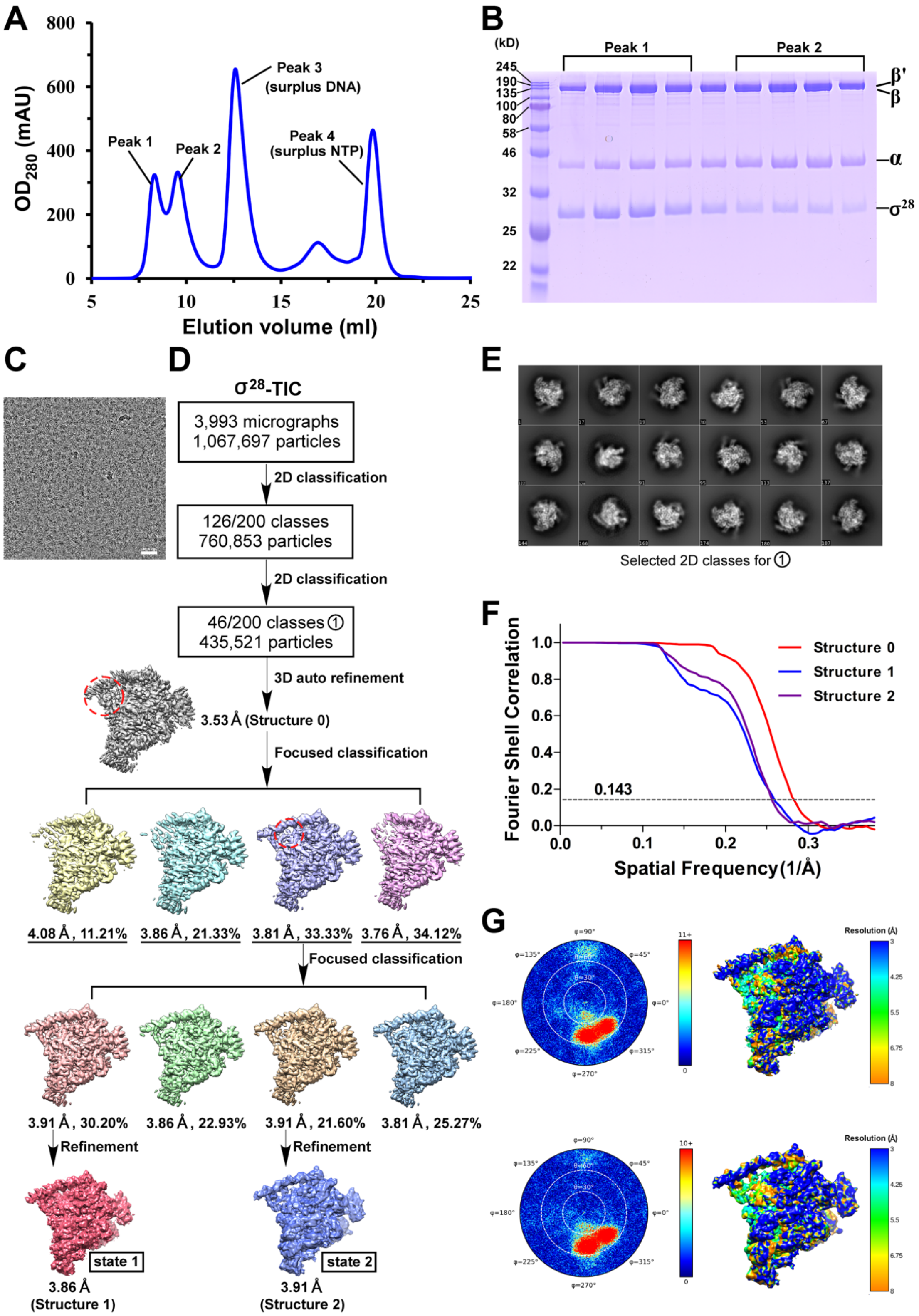

**Figure EV1. Isolation of the $\sigma^{28}$-TIC, cryo-EM images and data processing procedure for $\sigma^{28}$-TIC**

A. The size-exclusion chromatography profile of the $\sigma^{28}$-TIC is presented. Peak 2 is the target complex, while peaks 1, 3, and 4 are the aggregation form, surplus DNA, and surplus NTP, respectively.

B. The SDS-PAGE gel visualized the components and verified the presence of the complex.

C. A representative micrograph.

D. Flow chart of the cryo-EM image processing (see Method details).

E. Selected 2D classes for the structure 0.

F. Gold-standard Fourier Shell Correlations (FSCs) of the maps for structures 0-2.

G. Angular orientation distributions of the particles used in the final reconstruction and local resolution maps for the structure 1 (top) and structure 2 (bottom).

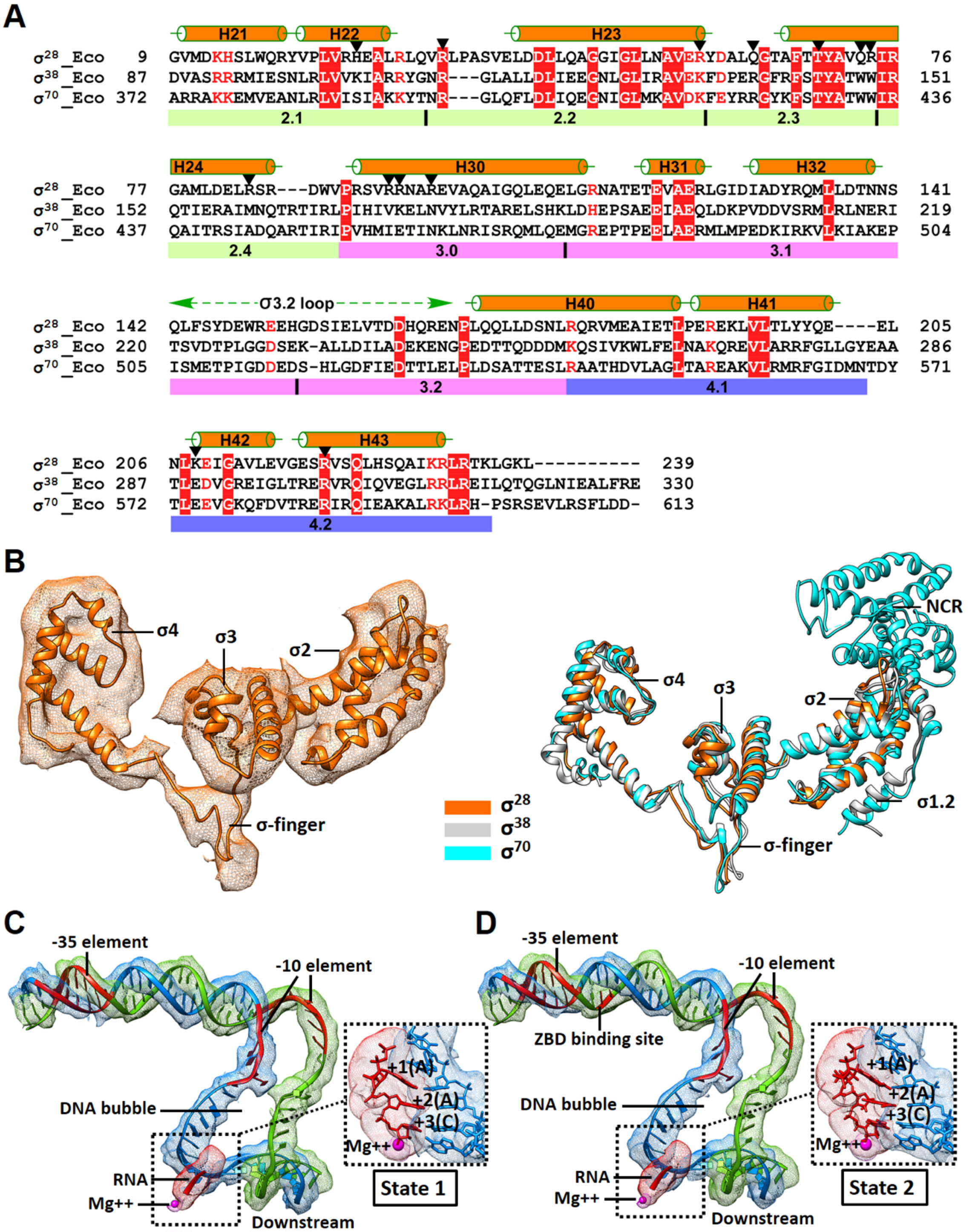


**Figure EV2. Alignments of the σ$^{28}$ factor with other factors and the nucleic acid structures in σ$^{28}$-TICs**

A. The sequence alignment of the $\sigma^{28}$ factor with the $\sigma^{38}$, $\sigma^{70}$ factor in *E. coli*. Sequences are presented with the one-letter amino acid codes. The second structure (α helices) and the conserved regions are labeled above and below the sequences, respectively. The black triangles are used to denote the promoter-recognition amino acids of $\sigma^{28}$ factor..

B. Overall structure of the *E. coli* $\sigma^{28}$ factor in the TICs. The transparent density map is contoured at 8 RMS. The *E. coli* $\sigma^{28}$ factor in the $\sigma^{28}$-TIC, the $\sigma^{38}$ factor in the $\sigma^{38}$-TIC (PDB ID 5IPL), and the $\sigma^{70}$ factor in the $\sigma^{70}$-TIC (PDB ID 4YLN) are superimposed.

C, D. The transparent split cryo-EM maps (contoured at 10 RMS) for the promoter DNA, the nascent RNA, and the active site Mg ion in the $\sigma^{28}$-TIC at the state 1 (**C**) and the state 2 (**D**) are shown, respectively. The red labeled on the promoter DNA are the important recognition regions. The color schemes for others are same as in Fig 1. The right insets are the zoom-in views of the DNA-RNA hybrid.

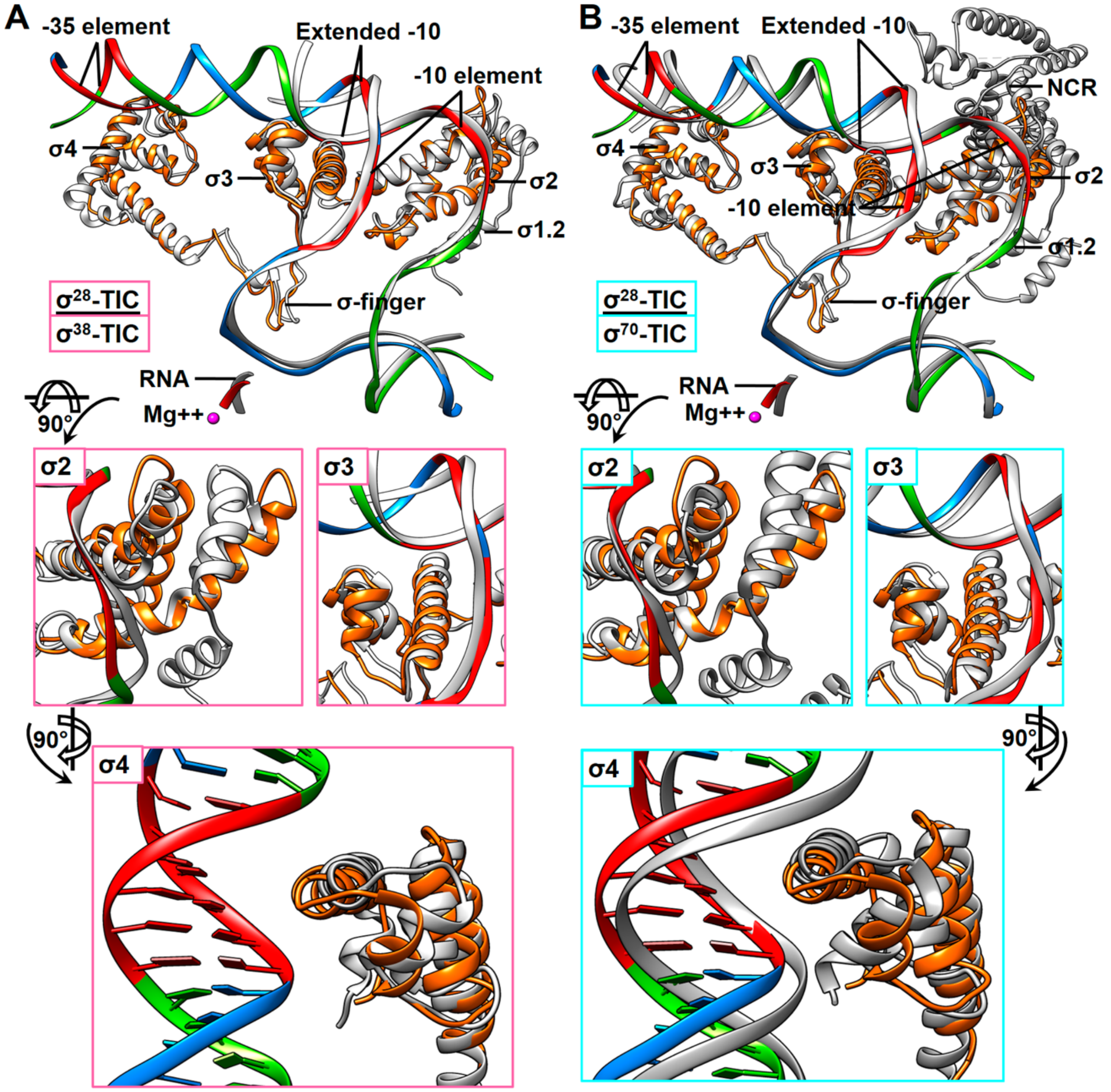


**Figure EV3. Comparisons among TICs on sigma factors and promoters**

A, B. Comparisons on sigma factors and the promoter DNAs between the state-1 $\sigma^{28}$-TIC and $\sigma^{38}$-TIC (**A**; gray, PDB ID 5IPL) or $\sigma^{70}$-TIC (**B**; gray, PDB ID 4YLN) are shown. The RNAP was omitted for clear representations. The conserved elements on the promoter DNA were labeled in red. Other color schemes are same as in Fig 1. The insets are zoom-in views of the comparisons at the σ2, σ3, and σ4 domains.

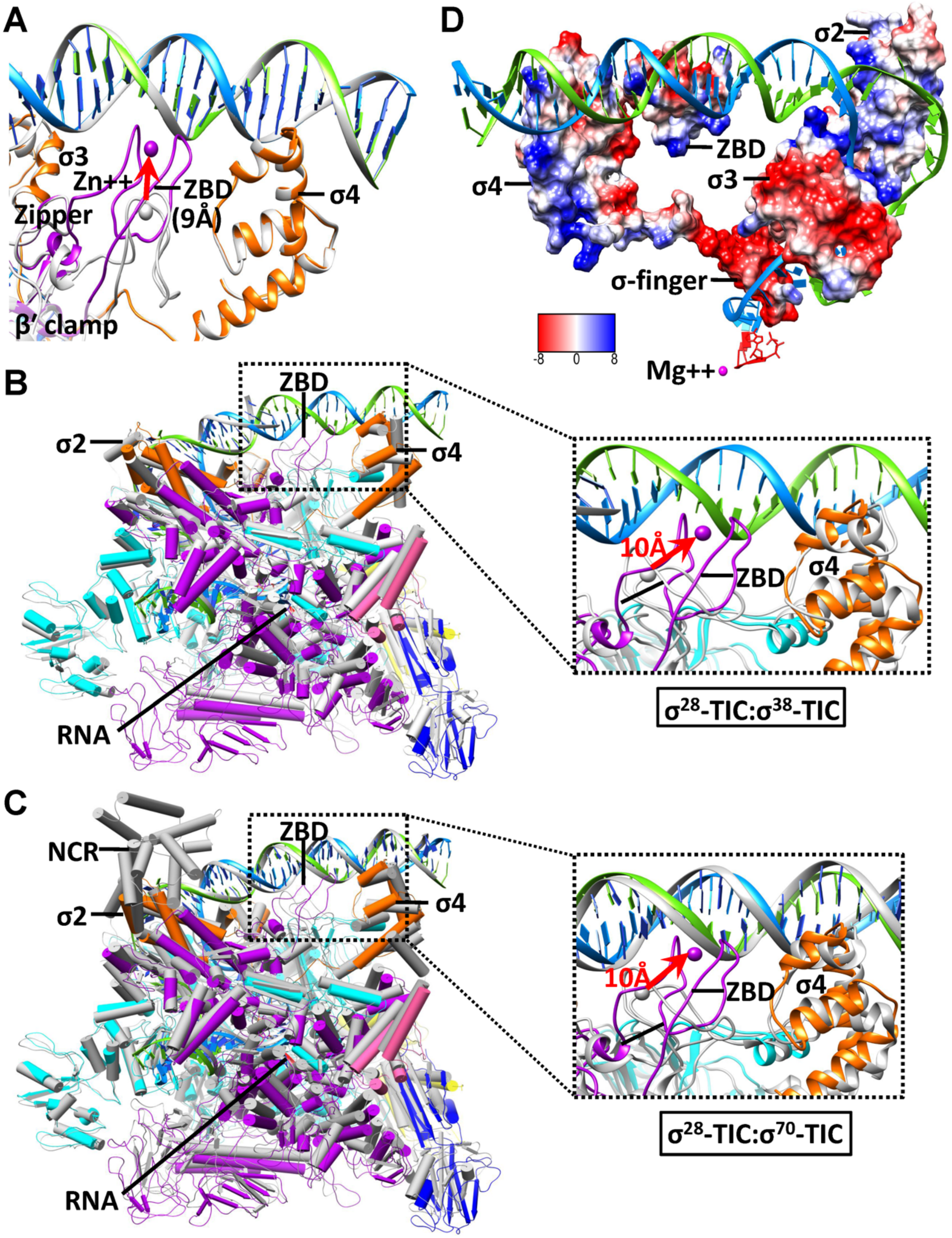


**Figure EV4. Comparisons among TICs on β′ ZBD**

A. Comparison between the two states of $\sigma^{28}$-TICs displays the shift of β′ ZBD (~9 Å) with its relocation to approach the upstream DNA. The color schemes for others are same as in Fig 1. The state-1 $\sigma^{28}$-TIC is colored in gray.

B, C. Comparisons between the state-2 $\sigma^{28}$-TIC and $\sigma^{38}$-TIC (**B**; PDB ID 5IPL) or $\sigma^{70}$-TIC (**C**; PDB ID 4YLN) via sigma factors are shown. The $\sigma^{38}$-TIC and $\sigma^{70}$-TIC are colored in gray. Other color schemes are same as in Fig 1. The insets are zoom-in views of the comparisons at β′ ZBD.

D. An electrostatic surface representation demonstrates the synergistic relation in recognizing and stabilizing the promoter DNA between β′ ZBD and the factor in the complex at the state 2.

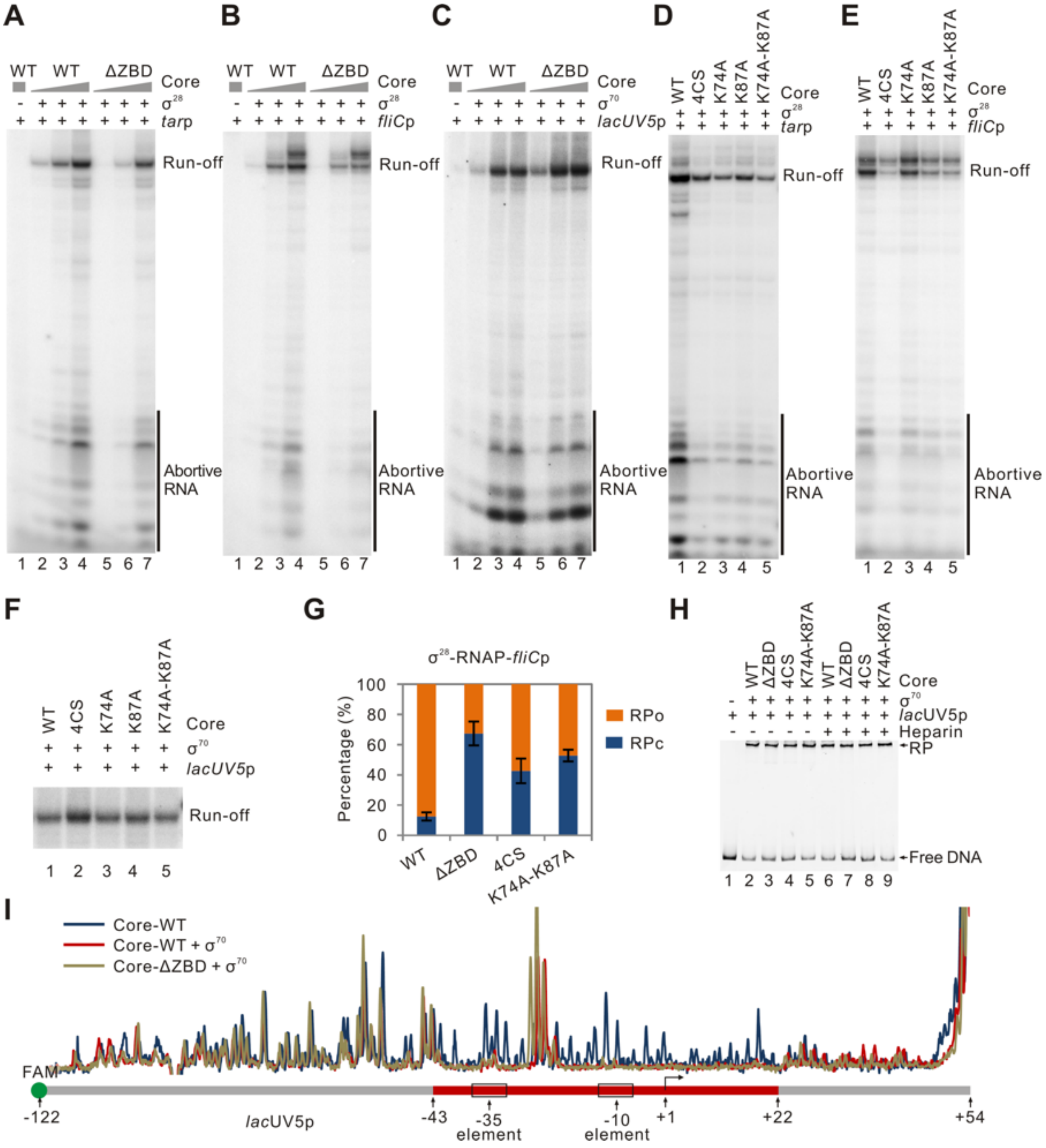


**Figure EV5. Role of β′ ZBD in promoter binding and transcription**

A-E. Full image of transcripts from wild-type (WT) or ZBD mutated RNAPs on different promoters as shown in Fig 3B-F. The amount of abortive transcripts showed a similar trend as run-off products for each sample.

F. Effects of deleting β′ ZBD on transcriptional activities of $\sigma^{70}$-RNAP on *lac*UV5p.

G. Percentages of RNAP-promoter open complex (RPo) and closed complex (RPc) in EMSA performed using $\sigma^{28}$-RNAP and *fliC*p. Data are mean ± SD from three replicates.

H. Bindings of $\sigma^{70}$-RNAP with *lac*UV5p as detected by EMSA.

I. DNase I footprinting of the *lac*UV5p complex with $\sigma^{70}$-RNAP. The promoter region protected by RNAP is indicated by red box at the bottom panel.

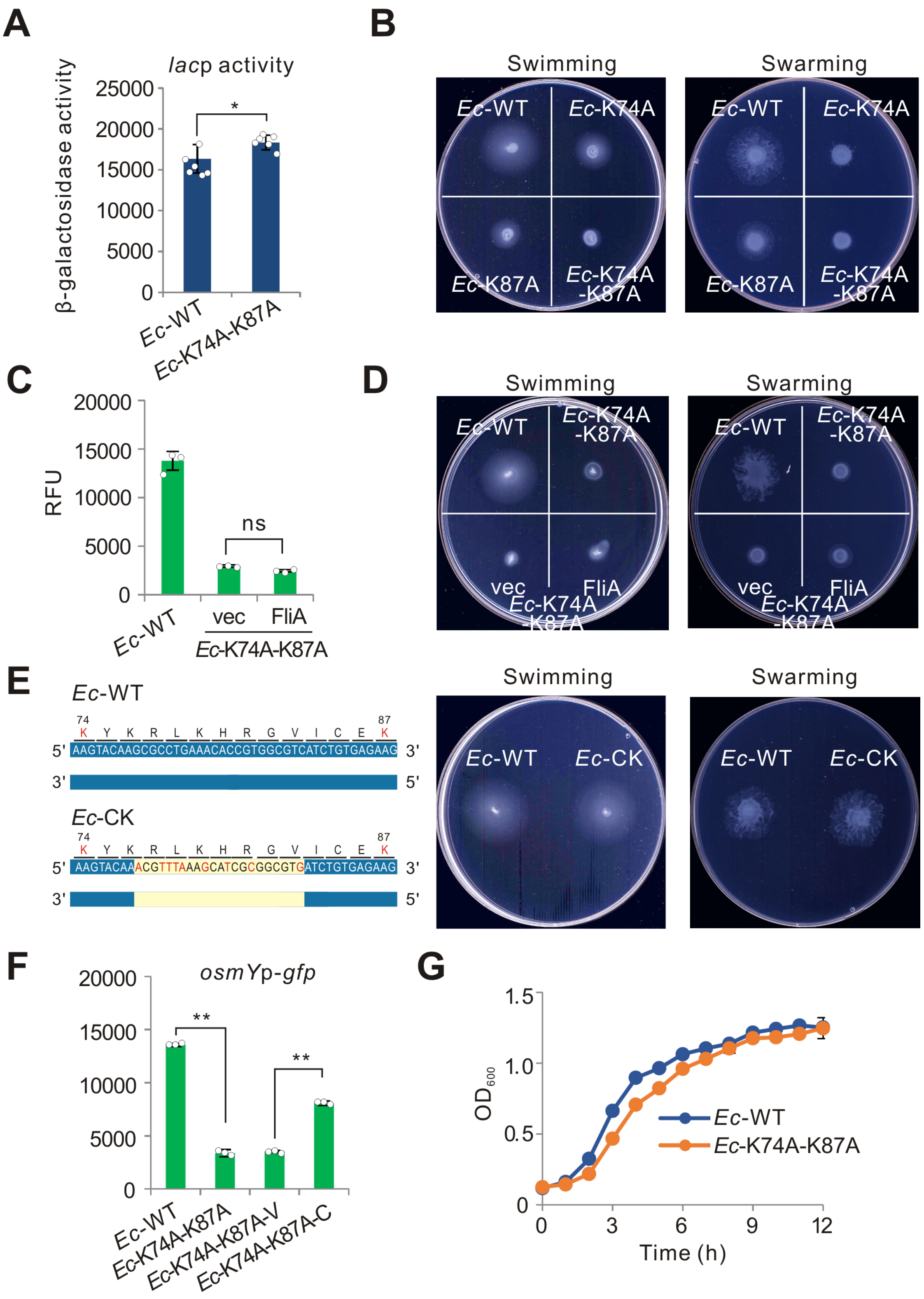


**Figure EV6. Role of β′ ZBD in controlling transcription *in vivo***

A. Activities of native *lac* promoter (*lac*p) in *Ec*-WT and *Ec*-K74A-K87A as measured by β-galactosidase test. Bacteria were grown to late logarithmic growth phase at 30 °C. Individual values of biological replicates (n=6) are shown as dots, and the mean ± SD values are displayed as error bars * $p<0.05$.

B. Motilities of *E. coli* wild-type and the β′ ZBD mutated strains.

C. Promoter activities of *fliC*p in *Ec*-WT strain and *Ec*-K74A-K87A strain carrying the control plasmid (vec) or with over-expression of FliA (FliA). L-arabinose was added at final concentration of 0.002%. Data are mean ± SD from three colonies.

D. Motilities of *Ec*-K74A-K87A strains with over-expression of wild-type *fliA* gene (FliA) or carrying the control plasmid (vec).

E. Motilities of *Ec*-WT and *Ec*-CK strains. Sequences of partial ZBD domain in *Ec*-WT and *Ec*-CK strains are shown in the left panel. Synonymous mutations introduced in the *Ec*-CK strain are indicated in yellow background.

F. Activities of *osmY*p in *Ec*-K74A-K87A strain with over-expression of wild-type *rpoC* gene (*Ec*-K74A-K87A-C) or carrying the empty plasmid (*Ec*-K74A-K87A-V). Data are mean ± SD from three colonies. ** $p<0.01$.

G. Growth curve of *Ec*-WT and *Ec*-K74A-K87A strains at 37 °C. Data are mean ± SD from three colonies.

**Table EV1. Statistics of cryo-EM data collection, 3D reconstruction and model building**

| **Data collection/processing** | **$\sigma^{28}$-TIC** | |
|---|---|---|
| Microscope | Krios | |
| Voltage (kV) | 300 | |
| Camera | Falcon III | |
| Camera mode | Counted | |
| Defocus range (μm) | −1.6 ~ −2.6 | |
| Exposure time (s) | 30 | |
| Dose rate ($e^-$/pixel/s) | 0.8 | |
| Magnified pixel size (Å) | 0.9 | |
| **Reconstruction (focused maps)** | **State 1** | **State 2** |
| Software | cisTEM | cisTEM |
| Symmetry | C1 | C1 |
| Particles refined | 43838 | 31354 |
| Resolution (auto-masked, Å) | 3.86 | 3.91 |
| Access code | EMD-20394 | EMD-20395 |
| **Model Statistics** | **State 1** | **State 2** |
| Number of residues (modeled) | 3567 | 3567 |
| Map CC (Chimera) | 0.8128 | 0.8073 |
| MolProbity score | 1.91 | 1.86 |
| All-atom Clashscore | 6.02 | 5.48 |
| Cβ deviations | 0 | 0 |
| Rotamer outliers | 0.27% | 0.24% |
| Ramachandran | | |
| Outliers | 0.38% | 0.35% |
| Favored | 88.94% | 89.29% |
| RMS deviations | | |
| Bond length | 0.009 | 0.008 |
| Bond angles | 1.160 | 1.102 |
| Access code | 6PMI | 6PMJ |

**Table EV2. Promoter sequences used in this study**

| Name | DNA sequence (5’ to 3’)* |
|---|---|
| *tar*p | ccggcccgatttcctcaattgaaatgaacccgatgatctgcgcatcgggttttttatttcaatttcgcggcgggtg gcatcagc**AATAAAG**tttccccctccttgc**CGATAAC**gagatc**A**acttgttttcaggaaggtgc cttatgattaaccgtatccgcgtagtcacgccggg |
| *fliC*p | cccgccgtttttaatagcgggaataaggggcagagaaaagagtatttcggcgactaacaaaaaatggctgtttt tgaaaaaaat**TCTAAAG**gttgttttacgacaga**CGATAAC**agggtt**G**acggcgattgagccgac gggtggaaacccaatacgtaatcaacgactgggcgcgg |
| *lac*UV5p | cccgccgacaggtttcccgactggaaagcgggcagtgagcgcaacgcaattaatgtgagttagctcactcatt aggcaccccaggc**TTTACA**ctttatgcttccggctcg**TATAAT**gtgtgg**A**attgtgagcggataac aatttcacacaggaaacagctatgaccatgggcgcgg |
| *osmY*p | cgccccgtcaatttcccttccttattagccgcttacggaatgttcttaaaacattcacttttgcttatgttttcgctgat atcc**CGAGCG**gtttcaaaattgtgatc**TATATT**taacaaa**G**tgatgacatttctgacggcgttaaata ccgttcaatgcgtagatatcgggcgcgg |
| *fliC*p-S[#] | ttttgaaaaaaaat**TCTAAAG**gttgttttacgacaga**CGATAAC**agggtt**G** |
| *fliD*p-S[#] | aaaaacaattaaa**CGTAAAC**tttgcgcaattcagac**CGATAAC**cccggt**A** |
| *fliA*p-S[#] | tccgattaaaaac**CCTGCAG**aaacggataatcatgc**CGATAAC**tcatat**A** |
| *motA*p-S[#] | ggcagtaaaaaga**CGTAAAC**tttcccagaatcctgc**CGATATT**atcccac**A** |
| *flgM*p-S[#] | aaacctgtaagct**GTTAAAG**attacccgtcccttgc**CGATAAA**taagca**A** |
| *flgK*p-S[#] | tctgttctgaata**ACTCAAG**tccggcgggtcgctgc**CGATAAT**actctgt**A** |
| *ycgR*p-S[#] | attcacgcatttg**TATTAAG**ttttgttaactgtgac**CGATAAA**ccaaag**A** |
| *tar*p-S[#] | gggtggcatcagc**AATAAAG**tttccccctccttgc**CGATAA**cgagatc**A** |

*Sequences proposed for −10 and −35 elements and transcriptional start site in each promoter were shown in **BOLD CAPITAL** letters.

[#] Promoter sequences used in conserved motif analysis shown in Fig 2C.

**Table EV3. Bacterial strains and plasmids used in this study**

| Name | application, or characters, or sequences | SOURCE |
|---|---|---|
| Strains | | |
| *E. coli* BL21(DE3) | Protein expression | Novagen |
| *E. coli* DH5α | Cloning construction | Shenzhen KT Life |
| *Ec*-WT | Wild-type *E. coli* strain MG1655 | Dr. Qingsheng Qi |
| *Ec*-K74A | *E. coli* MG1655 strain with mutation of K74A in *rpoC* gene | This study |
| *Ec*-K87A | *E. coli* MG1655 strain with mutation of K87A in *rpoC* gene | This study |
| *Ec*-K74A-K87A | *E. coli* MG1655 strain with mutations of both K74A and K87A in *rpoC* gene | This study |
| *Ec*-K74A-K87A-C | *Ec*-K74A-K87A carrying the pBAD-EcorpoC | This study |
| *Ec*-K74A-K87A-V | *Ec*-K74A-K87A carrying the pBAD22 | This study |
| *Ec*-CK | *E. coli* MG1655 strain carrying synonymous mutations shown in Fig EV 6E in *rpoC* gene | This study |
| Plasmids | | |
| pVS10-RNAP | Plasmid expressing *E. coli* RNAP core enzyme, β′ subunit with C-terminal $His_6$-tag. | (Belogurov et al, 2007) |
| pET21a-Ecoσ28 | Plasmid expressing *E. coli* $\sigma^{28}$, with C-terminal $His_6$-tag. | This paper |
| pET21a-Ecoσ28-K208A-R220A | Plasmid expressing K208A-R220A mutated *E. coli* $\sigma^{28}$, with C-terminal $His_6$-tag. | This paper |
| pET21a-Ecoσ28-R4m | Plasmid expressing σ mutated *E. coli* $\sigma^{28}$, with C-terminal His-tag. | This paper |
| pET28a-EcorpoD | Plasmid expressing *E. coli* $\sigma^{70}$, with N-terminal His-tag. | This paper |
| pET28a-MBP-tev-EcorpoS | Plasmid expressing *E. coli* $\sigma^{38}$, with N-terminal $His_6$-MBP-tev tag. | This paper |
| pVS10-RNAP-ΔZBD | pVS10-RNAP carrying deletion of ZBD region (amino acid 64 to 95) in *rpoC* gene | This paper |
| pVS10-RNAP-4CS | pVS10-RNAP carrying mutations of four cysteines in ZBD domain in *rpoC* gene | This paper |
| pVS10-RNAP-K74A | pVS10-RNAP carrying mutation of K74A in *rpoC* gene | This paper |

| | | |
|---|---|---|
| pVS10-RNAP-K87A | pVS10-RNAP carrying mutation of K87A in *rpoC* gene | This paper |
| pVS10-RNAP-K74A-K87A | pVS10-RNAP carrying mutations of both K74A and K87A in *rpoC* gene | This paper |
| pCas | Plasmid expressing spCas9 protein | (Jiang et al, 2015) |
| pTargetF | Plasmid for expressing sgRNA | (Jiang et al, 2015) |
| pTargetF-EcorpoC | Plasmid expressing sgRNA targeting to ZBD region of *rpoC* gene in *E. coli* | This paper |
| pBAD22 | Arabinose inducible gene expression plasmid | (Guzman et al, 1995) |
| pBAD-EcorpoC | pBAD22 expressing wild-type RNAP β′ subunit from *E. coli* | This paper |
| pBAD-FliA | pBAD22 expressing wild-type FliA protein from *E. coli* | This paper |
| pGT | Promoter-GFP fusion plasmid | This paper |
| pGT-*tar*p | pGT carrying fusion of *tar*p-GFP | This paper |
| pGT-*fliC*p | pGT carrying fusion of *fliC*p-GFP | This paper |
| pGT-*osmY*p | pGT carrying fusion of *osmY*p-GFP | This paper |
| pRK793 | Plasmid expressing $His_6$-TEV protease | (Kapust et al, 2001) |

#### Table EV4. Oligonucleotides used in this study

| Application and sequence (5'-3') | Source |
|---|---|
| Sigma28 forward primer<br>GGAATTCCATATGAATTCACTCTATACCGCTGAAGG | IDT |
| Sigma28 reverse primer<br>GCTAATCTCGAGTAACTTACCCAGTTTAGTGCGTAAC | IDT |
| Specific promoter non-template DNA<br>AGCAATAAAGTTTCCTTCCTCCTTGCCGATAACGAGAT<br>CAACTTGTTTGCGGCG | IDT |
| Specific promoter template DNA<br>CGCCGCAAACAAGTTGTAGAGCTTATCGGCAAGGAGG<br>AAGGAAACTTTATTGCT | IDT |
| Sigma70 forward primer<br>ACACGCTAGCATGGAGCAAAACCCGCAGTCAC | TIANYI HUIYUAN, China |
| Sigma70 reverse primer<br>ACCGAAGCTTTTAATCGTCCAGGAAGCTACGCAG | TIANYI HUIYUAN, China |
| Sigma38 forward primer<br>GAACCTGTACTTCCAATCCATGAGTCAGAATACGCTGAA<br>AGTTCATG | TIANYI HUIYUAN, China |
| Sigma38 reverse primer<br>CTTCCTTTCGGGCTTTGTTACTCGCGGAACAGCGCTTC<br>GATATTCAGCCC | TIANYI HUIYUAN, China |
| pET-MBP forward primer<br>TAACAAAGCCCGAAAGGAAGCTGA | TIANYI HUIYUAN, China |
| pET-MBP reverse primer<br>GGATTGGAAGTACAGGTTCTCTC | TIANYI HUIYUAN, China |
| Sigma28-K208A-R220A forward primer<br>GAGATTGGCGCGGTGCTGGAGGTCGGGGAATCGGCG<br>GTCAGTCAGTTACACAGCCAGGC | TIANYI HUIYUAN, China |
| Sigma28-K208A-R220A reverse primer<br>CAGCACCGCGCCAATCTCGGCGAGATTCAGCTCTTCCT<br>GGTAATAGAGGGTTAATACC | TIANYI HUIYUAN, China |
| Sigma28-R4m forward primer<br>CACCACCACTGAGATCCGGCTGCTAACAAAGCCCGAAA<br>GGAAGCTGA | TIANYI HUIYUAN, China |
| Sigma28-R4m reverse primer<br>GTCGCAGAATCCAGCGGGTTTTCTCGCTGATGATCATC<br>AGTAACCAG | TIANYI HUIYUAN, China |
| Sigma70-R4 forward primer<br>CCGCTGGATTCTGCGACCACCGAAAG | TIANYI HUIYUAN, China |
| Sigma70-R4 reverse primer<br>TTCCTTTCGGGCTTTGTTAATCGTCCAGGAAGCTACGC<br>AG | TIANYI HUIYUAN, China |
| EcorpoC-SbfI forward primer<br>CTTACAGCCTGGTTACTCAGCAGC | TIANYI HUIYUAN, China |
| EcorpoC-HindIII reverse primer<br>CAGTACCAGGTCAAAACGGTTACC | TIANYI HUIYUAN, China |
| pVS10-K74A forward primer<br>CCTGTGCGGTGCGTACAAGCGCCTGAAACACCGTGGC | TIANYI HUIYUAN, China |
| pVS10-K74A reverse primer<br>GGCGCTTGTACGCACCGCACAGGCACTCGTAATCT | TIANYI HUIYUAN, China |

| | |
|---|---|
| pVS10-K87A forward primer CATCTGTGAGGCGTGCGGCGTTGAAGTGACCCAGACTAAAG | TIANYI HUIYUAN, China |
| pVS10-K87A reverse primer CAACGCCGCACGCCTCACAGATGACGCCACGGTG | TIANYI HUIYUAN, China |
| pVS10-K74-K87A forward primer CACCGTGGCGTCATCTGTGAGGCGTGCGGCGTTGAAGTGACCCAGACTAAAG | TIANYI HUIYUAN, China |
| pVS10-K74-K87A reverse primer ACAGATGACGCCACGGTGTTTCAGGCGCTTGTACGCACCGCACAGGCACTCGTAATCT | TIANYI HUIYUAN, China |
| pVS10-ΔZBD forward primer TATCTTTGGGGGTAGCAAAGTACGCCGTGAGCGTATGGGCC | TIANYI HUIYUAN, China |
| pVS10-ΔZBD reverse primer GGCGTACTTTGCTACCCCCAAAGATACGGGCGCAGAAAAGGCCGT | TIANYI HUIYUAN, China |
| pVS10-4CS forward primer AAGCGCCTGAAACACCGTGGCGTCATCAGCGAGAAGAGCGGCGTTGAAGTGACCCAGAC | TIANYI HUIYUAN, China |
| pVS10-4CS reverse primer GGTGTTTCAGGCGCTTGTACTTACCGCTCAGGCTCTCGTAATCTTTTACCGGCCCAAAG | TIANYI HUIYUAN, China |
| lacUV5p forward primer (5’-FAM labeled) CCCGCCGACAGGTTTCCCGACTGGA | TIANYI HUIYUAN, China |
| lacUV5p reverse primer 1 GTTATCCGCTCACAATTCCACACATTATACGAGCCGGAAGCATAAAGTGTAAAG | TIANYI HUIYUAN, China |
| lacUV5p reverse primer 2 CCGCGCCCATGGTCATAGCTGTTTCCTGTGTGAAATTGTTATCCGCTCACAATTCCAC | TIANYI HUIYUAN, China |
| fliCp forward primer (5’-FAM labeled) CCCGCCGTTTTTAATAGCGGGAATAAGGGGCAGAG | TIANYI HUIYUAN, China |
| fliCp reverse primer CCGCGCCCAGTCGTTGATTACGTATTGGGTTTC | TIANYI HUIYUAN, China |
| osmYp forward primer CGCCCCGTCAATTTCCCTTCCTTATTAGCCG | TIANYI HUIYUAN, China |
| osmYp reverse primer CCGCGCCCGATATCTACGCATTGAACGGTATTTAACG | TIANYI HUIYUAN, China |
| tarp forward primer CCGGCCCGATTTCCTCAATTGAAATGAACCCGATG | TIANYI HUIYUAN, China |
| tarp reverse primer CCCGGCGTGACTACGCGGATACGGTTAATC | TIANYI HUIYUAN, China |
| tarp-M1 forward primer GTGGCATCAGCTTGACATTTCCCCCCTCCTTGCCGATAACG | TIANYI HUIYUAN, China |
| tarp-M1 reverse primer TGTCAATGCTGATGCCACCCGCCGCGAAAT | TIANYI HUIYUAN, China |
| tarp-M forward primer ATAACGAGATCAACTTGTTTTCAGGAAGG | TIANYI HUIYUAN, China |
| tarp-M2 reverse primer AACAAGTTGATCTCGTTATAGGCAAGGAGGGGGGAAACTTTATTGCTG | TIANYI HUIYUAN, China |

| | |
|---|---|
| tarp-M3 reverse primer AACAAGTTGATCTCGTTATTGGCAAGGAGGGGGGAAACTTTATTGCTG | TIANYI HUIYUAN, China |
| tarp-M4 reverse primer AACAAGTTGATCTCGTTATGGGCAAGGAGGGGGGAAACTTTATTGCTG | TIANYI HUIYUAN, China |
| tarp-M5 reverse primer AACAAGTTGATCTCGTTATCCGCAAGGAGGGGGGAAACTTTATTGCTG | TIANYI HUIYUAN, China |
| tarp-M6 reverse primer AACAAGTTGATCTCGTTATCTGCAAGGAGGGGGGAAACTTTATTGCTG | TIANYI HUIYUAN, China |
| tarp-M7 reverse primer AACAAGTTGATCTCGTTATCAGCAAGGAGGGGGGAAACTTTATTGCTG | TIANYI HUIYUAN, China |
| pTargetF-EcorpoC forward primer GTATAATACTAGTAGATGACGCCACGGTGTTTCGTTTTAGAGCTAGAAATAGCAAGTT | TIANYI HUIYUAN, China |
| pTargetF-EcorpoC reverse primer GCTCTAAAACGAAACACCGTGGCGTCATCTACTAGTATTATACCTAGGACTGAGCTAG | TIANYI HUIYUAN, China |
| *Ec*-K74A forward primer GCGTACAAACGTTTAAAGCATCGCGGCGTGATCTGTGAGAAGTGCGGCGT | TIANYI HUIYUAN, China |
| *Ec*-K74A reverse primer CGCCGCGATGCTTTAAACGTTTGTACGCACCGCACAGGCACTCGTAATCT | TIANYI HUIYUAN, China |
| *Ec*-K87A forward primer CGTTTAAAGCATCGCGGCGTGATCTGTGAGGCGTGCGGCGTTGAAGTGACCCAGACTA | TIANYI HUIYUAN, China |
| *Ec*-K87A reverse primer CGCCGCGATGCTTTAAACGTTTGTACTTACCGCACAGGCACTCG | TIANYI HUIYUAN, China |
| pBAD22 forward primer AAGCTTGGCTGTTTTGGCGG | TIANYI HUIYUAN, China |
| pBAD22 reverse primer CATGGTGAATTCCTCCTGCTAG | TIANYI HUIYUAN, China |
| EcorpoC-BAD forward primer CAGGAGGAATTCACCATGAAAGATTTATTAAAGTTTCTGAAAGCG | TIANYI HUIYUAN, China |
| EcorpoC-BAD reverse primer GCCAAAACAGCCAAGCTTACTCGAGCTCGTTATCAGAACCG | TIANYI HUIYUAN, China |
| FliA-BAD forward primer CAGGAGGAATTCACCATGAATTCACTCTATACCGCTGAAG | TIANYI HUIYUAN, China |
| FliA-BAD reverse primer GCCAAAACAGCCAAGCTTTTATAACTTACCCAGTTTAGTGCG | TIANYI HUIYUAN, China |
| pGT forward primer CTAAGTCGACTCTAGAGAAGGAGATATACATATGGCTAGCAAAGGAGAAGAACTTTTC | TIANYI HUIYUAN, China |
| pGT reverse primer GGCCGTGACGTGACTAGTAAAAAAGGG | TIANYI HUIYUAN, China |

| | |
|---|---|
| pGT-fliCp forward primer CTAGTCACGTCACGGCCTTTTTAATAGCGGGAATAAGGGGCAGAG | TIANYI HUIYUAN, China |
| pGT-fliCp reverse primer TCTCTAGAGTCGACTTAGAGTCGTTGATTACGTATTGGGTTTC | TIANYI HUIYUAN, China |
| pGT-tarp forward primer CTAGTCACGTCACGGCCCCCGATTTCCTCAATTGAAATGAACC | TIANYI HUIYUAN, China |
| pGT-tarp reverse primer TCTCTAGAGTCGACTTAGGCGTGACTACGCGGATACGGTTAATC | TIANYI HUIYUAN, China |
| pGT-osmYp forward primer CTAGTCACGTCACGGCCGTCAATTTCCCTTCCTTATTAGCCG | TIANYI HUIYUAN, China |
| pGT-osmYp reverse primer TCTCTAGAGTCGACTTAGGATATCTACGCATTGAACGGTATTTAACG | TIANYI HUIYUAN, China |
| 16s rRNA qPCR forward primer CAGCCACACTGGAACTGAGA | TIANYI HUIYUAN, China |
| 16s rRNA qPCR reverse primer GTTAGCCGGTGCTTCTTCTG | TIANYI HUIYUAN, China |
| *fliC* qPCR forward primer AGGTTGGCGCAAATGATAAC | TIANYI HUIYUAN, China |
| f*liC* qPCR reverse primer AGTGGCTGCTTCCGTAGAAA | TIANYI HUIYUAN, China |
| *fliD* qPCR forward primer AAATACACCGCCGTAGATGC | TIANYI HUIYUAN, China |
| *fliD* qPCR reverse primer TCAACTGCGTCTGAATCGTC | TIANYI HUIYUAN, China |
| *motA* qPCR forward primer ATGCAGTGCGTCAAAGTCAC | TIANYI HUIYUAN, China |
| *motA* qPCR reverse primer GCACATGCTCTTCCAGTTCA | TIANYI HUIYUAN, China |
| *tar* qPCR forward primer TCACCAATAAACCGCAAACA | TIANYI HUIYUAN, China |
| *tar* qPCR reverse primer TTGTTCAGCAATTCGCAGTC | TIANYI HUIYUAN, China |
| *fliA* qPCR forward primer CCGCTGAAGGTGTAATGGAT | TIANYI HUIYUAN, China |
| *fliA* qPCR reverse primer TCCTTGTAGGGCGTCATAGC | TIANYI HUIYUAN, China |
| *flhD* qPCR forward primer CTCCGAGTTGCTGAAACACA | TIANYI HUIYUAN, China |
| *flhD* qPCR reverse primer AACGTTGTCGCCATTTCTTC | TIANYI HUIYUAN, China |
